# Neddylation Regulates Mitochondrial Dynamics and Turnover in the Adult Heart

**DOI:** 10.64898/2025.12.03.692150

**Authors:** Jianqiu Zou, Wenjuan Wang, Chen Qu, Lingxian Zhang, Josue Zambrano-Carrasco, Yilang Li, Yi Lu, Hongyi Zhou, Kunzhe Dong, Tohru Fukai, Masuko Ushio-Fukai, Dawn Bowles, Michelle Mendiola Pla, Ryan Gross, Weiqin Chen, Jiliang Zhou, Jie Li, Huabo Su

## Abstract

**BACKGROUND:** Disruption of mitochondrial homeostasis drives cardiomyopathy and heart failure, yet upstream regulatory mechanisms remain poorly defined. Neddylation, a reversible post-translational conjugation of the ubiquitin-like protein NEDD8 by E1/E2/E3 enzymes, is essential for cardiac morphogenesis, but its role in the adult heart is unknown.

**METHODS:** We assessed the relevance of neddylation to human cardiac disease by gene set enrichment analysis of ischemic (ICM) and non-ischemic cardiomyopathy (NICM) datasets and by immunoblotting and qPCR of ventricular tissue from patients with ICM or dilated cardiomyopathy (DCM). In adult mice, we induced cardiomyocyte-restricted deletion of the NEDD8-activating enzyme 1 (NAE1) by tamoxifen injection and monitored cardiac function at baseline and after transverse aortic constriction (TAC). Bulk RNA-seq 4 weeks post-tamoxifen was combined with bioenergetic, biochemical, and ultrastructural analyses. To assess mitochondrial dynamics, we generated NAE1/MFN2 and NAE1/DRP1 double-knockout mice. Cullin activity, mitochondrial ubiquitination, and mitophagy were measured in hearts and cultured cardiomyocytes.

**RESULTS:** Neddylation pathways were dysregulated in human ICM and NICM datasets and in failing ICM/DCM myocardium. Cardiomyocyte-specific NAE1 deletion caused systolic dysfunction and heart failure by 10 weeks post-tamoxifen, culminating in premature death and exacerbating TAC-induced pressure-overload heart failure. At 4 weeks, NAE1 loss repressed metabolic and mitochondrial bioenergetic programs, reduced ATP production, and impaired respiration. Electron microscopy revealed elongated mitochondria and accumulated mitophagic vesicles, with dysregulation of DRP1, MFN2, PINK1, LC3-II, and p62. DRP1/NAE1 co-deletion accelerated systolic failure relative to either single knockout, whereas MFN2/NAE1 co-deletion did not alter early disease progression, implicating pathogenic mitochondrial hyperfusion. Genetic NAE1 depletion in vivo and pharmacologic NAE1 inhibition in vitro impaired mitophagic vesicle formation and flux, inactivated cullin scaffold proteins, reduced mitochondrial ubiquitination, and blunted mitophagic clearance.

**CONCLUSIONS:** Cardiac neddylation preserves adult heart function by coordinating mitochondrial fusion-fission dynamics and sustaining cullin-dependent ubiquitination and turnover of damaged mitochondria. These findings identify neddylation as a key regulator of mitochondrial quality control and link its disruption to human cardiomyopathy. Therapeutically, targeting the neddylation-cullin axis may limit mitochondrial dysfunction, enhance mitophagy, and improve energetic reserve in failing hearts, while neddylation signatures in patient myocardium may help guide stratification and precision therapy for cardiomyopathy.

**Clinical Perspective:** *What Is New?:* • Demonstrates for the first time that the NEDD8-activating enzyme (NAE1)driven neddylation pathway is indispensable for maintaining mitochondrial quality control in the adult heart. • Links loss of neddylation to mitochondrial hyperfusion, impaired mitophagy, and rapid progression to heart failure. • Reveals that neddylation promotes cullin-RING ligase-mediated ubiquitination of damaged mitochondria, coupling mitochondrial dynamics with turnover.

*What Are the Clinical Implications?:* • Restoring or enhancing cardiac neddylation may represent a novel therapeutic avenue for cardiomyopathies characterized by mitochondrial dysfunction. • Pharmacologic agents that bolster DRP1-dependent fission or activate cullin neddylation could potentially normalize mitochondrial dynamics and improve myocardial energetics. • Conversely, systemic neddylation inhibitors now in oncology trials warrant careful cardiac monitoring, as they may precipitate mitochondrial injury and heart failure. • Circulating or tissue markers of neddylation might help stratify patients at heightened risk for mitochondrial-driven cardiac disease and guide precision therapy.

## INTRODUCTION

Cardiovascular diseases represent the paramount cause of mortality globally, with cardiomyopathies contributing substantially to the aggravation of cardiac ailments by precipitating arrhythmias, heart failure, myocardial infarction, and other cardiovascular complications^1,2^. Metabolic disorders have been implicated in predominant variants of cardiomyopathy, encompassing dilated cardiomyopathy (DCM) and hypertrophic cardiomyopathy (HCM)^3,4^. Given the crucial role of mitochondria as the principal energy-generating organelle within cardiomyocytes (CMs)^5^, the maintenance of mitochondrial integrity is vital for cardiac wellness.

Mitochondria are dynamic organelles, with their remodeling regulated through biogenesis, dynamics (fusion and fission), and turnover (mitophagy)^6,7^, orchestrated by a family of GTPases, including Drp1^8^ for fission and Mfn1/2^9,10^ for fusion. These proteins, pivotal in sustaining mitochondrial dynamics, are profusely expressed in adult hearts^11^. Perturbations in mitochondrial dynamics compromise cellular respiration, provoke oxidative stress, and may lead to cardiomyopathy and heart failure, while restoration of these processes can ameliorate cardiac diseases^12^. Nonetheless, the molecular mechanism overseeing mitochondrial dynamics in CMs remain incompletely elucidated.

Mitophagy, the selective autophagic elimination of damaged mitochondria, is essential for maintaining mitochondrial fidelity^13^. Impairments in mitophagy result in the accumulation of dysfunctional mitochondria, culminating in CM lethality and cardiac dysfunction^14^. This process is initiated by the ubiquitination of outer mitochondrial membrane proteins by the Ub ligase Parkin, which aids in the recruitment of mitophagy receptors (e.g. p62 and NBR1) and the ensuing removal of compromised mitochondria via LC3-incorporated autophagosome^15,16^. Despite Parkin’s important role in mitophagy, its ablation in murine models induces only minimal mitochondrial anomalies and does not markedly impair fundamental cardiac performance^17–19^, suggesting the presence of alternative compensatory mechanisms involving other Ub ligases, whose identities and roles in CMs are yet to be determined.

NEDD8 (neural precursor cell expressed, developmentally downregulated 8) is an Ub-like protein that modifies target proteins through neddylation, mediated by NEDD8 specific E1 (a heterodimer of NAE1 and UBA3)-E2 (UBC12)-E3 enzymes and reversible by NEDD8 proteases such as the COP9 signalosome (CSN) and NEDP1 (Fig. 1A)^20^. Neddylation affects various cellular processes, and disruptions in this pathway can lead to a range of diseases, such as developmental defects, tumorigenesis, metabolic disease, and neurodegenerative disorders^21,22^. Currently, the biological roles of neddylation are largely understood through its effects on Cullins (Cullin 1-7), which are the most thoroughly investigated targets of NEDD8. The neddylation process activates Cullins, leading to the formation and activation of Cullin-RING ubiquitin ligases (CRLs)^23^. These complexes are responsible for the degradation of approximately 20% of cellular proteins, encompassing numerous critical regulators of the cell cycle^24^. We have previously demonstrated the critical role of neddylation in cardiac development and maturation via mediating CM proliferation and metabolism^25–29^. However, the significance of neddylation in postmitotic organs and its regulatory mechanisms in adult CMs, particularly through non-Cullin proteins, remains poorly understood.

**Figure 1.**
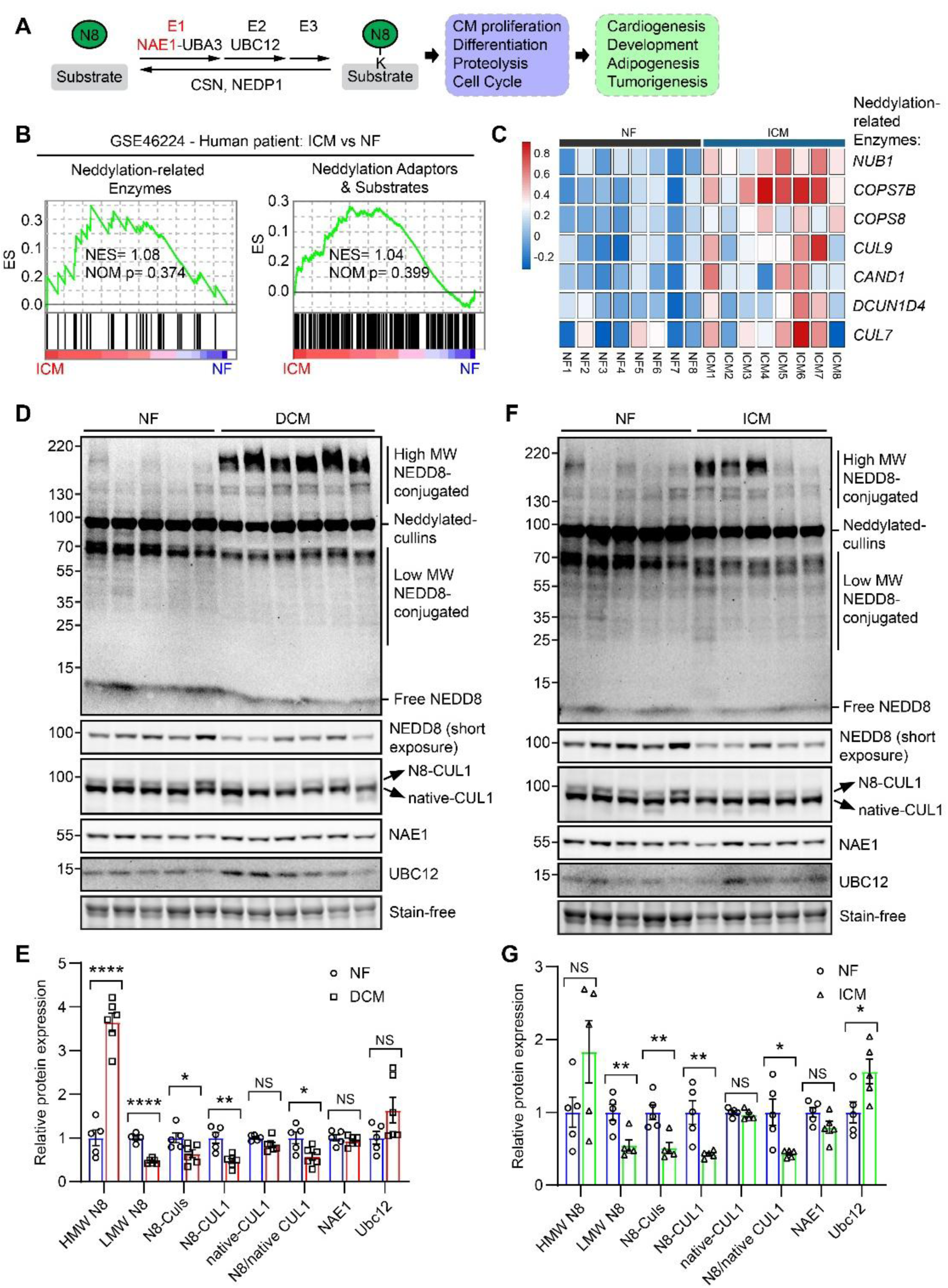
Implications of neddylation in cardiomyopathies and heart failure in human patients. **A**, neddylation pathway and its cellular and biological functions. **B**, Gene set enrichment analysis (GSEA) with deep RNA-seq data (GSE46224) of ischemic cardiomyopathy (ICM, n=8) patient group vs non-failing group (NF, n=8). Neddylation-related enzymes and neddylation adaptors and substrates as well as their specific GSEA plots. NES, normalized enrichment score; NOM p, nominal p-value. **C**, heat map showing representing dysregulated neddylation-related enzymes. **D** & **E**, western blot and quantification of dilated cardiomyopathy (DCM, n=6) patient hearts vs non-failing (NF, n=5) hearts. **F** & **G**, western blot and quantification of ischemic cardiomyopathy (ICM, n=5) patient hearts vs non-failing (NF, n=5) hearts. Statistical tests in panel **E** & **G** were nested t-test. *, 0.01<P< 0.05; **, 0.001<P< 0.01; ***, 0.0001<P< 0.001; ****, P< 0.0001; NS, not significant.

Here, we report the findings of dysregulated neddylation in both human patients and in diseased animal models, and how neddylation controls cardiac mitochondrial dynamics and turnover in multiple transgenic mouse models.

## RESULTS

### Dysregulation of neddylation in failing hearts with different etiologies

To investigate the involvement of neddylation in human patients with cardiomyopathies, we first reanalyzed the publicly available transcriptome datasets from published resources. We performed gene sets enrichment analysis (GSEA) on transcriptomes including 8 non-failing heart healthy individuals (NF) and 8 ischemic cardiomyopathy patients (ICM) (GSE46224, Gene Expression Omnibus)^30^. It was shown that neddylation-related enzymes (NES= 1.08) and neddylation adaptors and substrates (NES= 1.04) are both upregulated in the ICM group (Fig. 1B). Specifically, neddylation enzymes such as NUB1, CAND1 and some COPSs (COP9 Signalosome Complex Subunits) and CULs (cullins) showed upregulated expression in ICM samples (Fig. 1C). Meanwhile, non-ischemic cardiomyopathy (NICM) patient hearts from the same source (GSE46224) also showed upregulated neddylation enzymes and adaptors/substrates (Sup. Fig. 1A-C). In addition, in another set of online data (GSE48166, Gene Expression Omnibus), the sets of neddylation adaptors and substrates were upregulated in ICM (n=15) compared to NF (n=15) patient hearts, while the neddylation enzymes were downregulated (Sup. Fig. 1D-F). These data together suggested a dysregulated neddylation pathway in cardiomyopathy patients.

To further validate the implication of neddylation in cardiomyopathy patient hearts, we analyzed heart samples from healthy non-failing individuals (n=5), dilated cardiomyopathy patients (n=6), and ischemic cardiomyopathy patients (n=5) (Fig. 1DG). It was shown that higher molecular weight neddylated proteins (>130 kDa) were upregulated significantly, while lower molecular weight neddylated proteins (<70 kDa), neddylated cullins and neddylated CUL1 were significantly downregulated in both DCM (Fig. 1D-E) and ICM (Fig. 1F-G) samples. In addition, qPCR analysis revealed that in ICM samples, gene expression level of UBC12, NEDD8, CSN8, and CSN5 were significantly upregulated (Sup. Fig. 1G). These findings further validated the dysregulated neddylation pathway in patient hearts with cardiomyopathies, suggesting a critical role of neddylation in the progression of human heart diseases.

### Neddylation is required for the normal function of adult hearts

NAE1 is the regulatory subunit of the only NEDD8 E1 enzyme^23^ (Fig. 1A). To investigate the role of neddylation in post-mitotic hearts, we created tamoxifen (TAM)inducible cardiac restricted NAE1 knockout (NAE1KO) mice by crossing the αMHC^MerCreMer^ (MCM) mice^31^ with *Nae1*^flox/flox^ mice^32^ (Fig. 2A). We first titrated the doses of TAM to ensure efficient *Nae1* deletion without introducing confounding cardiotoxicity induced by MCM^31^. Administration of high dose of TAM (100 mg/kg/day for 5 days) via intraperitoneal injections led to the rapid development of heart failure in a week and early lethality in NAE1KO mice even compared with MCM, whereas administration of medium dose of TAM (50 mg/kg/day for 5 days) caused severe cardiomyopathy in both NAE1KO and MCM mice when compared with *Nae1*^flox/flox^ mice (Sup. Fig. 2A-2G). In contrast, delivery of TAM at 20 mg/kg/day for 10 days with a 2day interval after the first 5 injections did not cause any discernible cardiac dysfunction in *Nae1*^flox/flox^, MCM and NAE1KO mice following the TAM injections (Fig. 2B-D). This regimen was sufficient to effectively delete *Nae1* gene in the NAE1KO hearts, which consequently lead to decreased neddylated CUL2, neddylated CUL4A, and total neddylated proteins (Fig. 2E-F). Hence, we used this regimen to delete *Nae1* in the heart and probe its impact on cardiac function thereafter.

**Figure 2.**
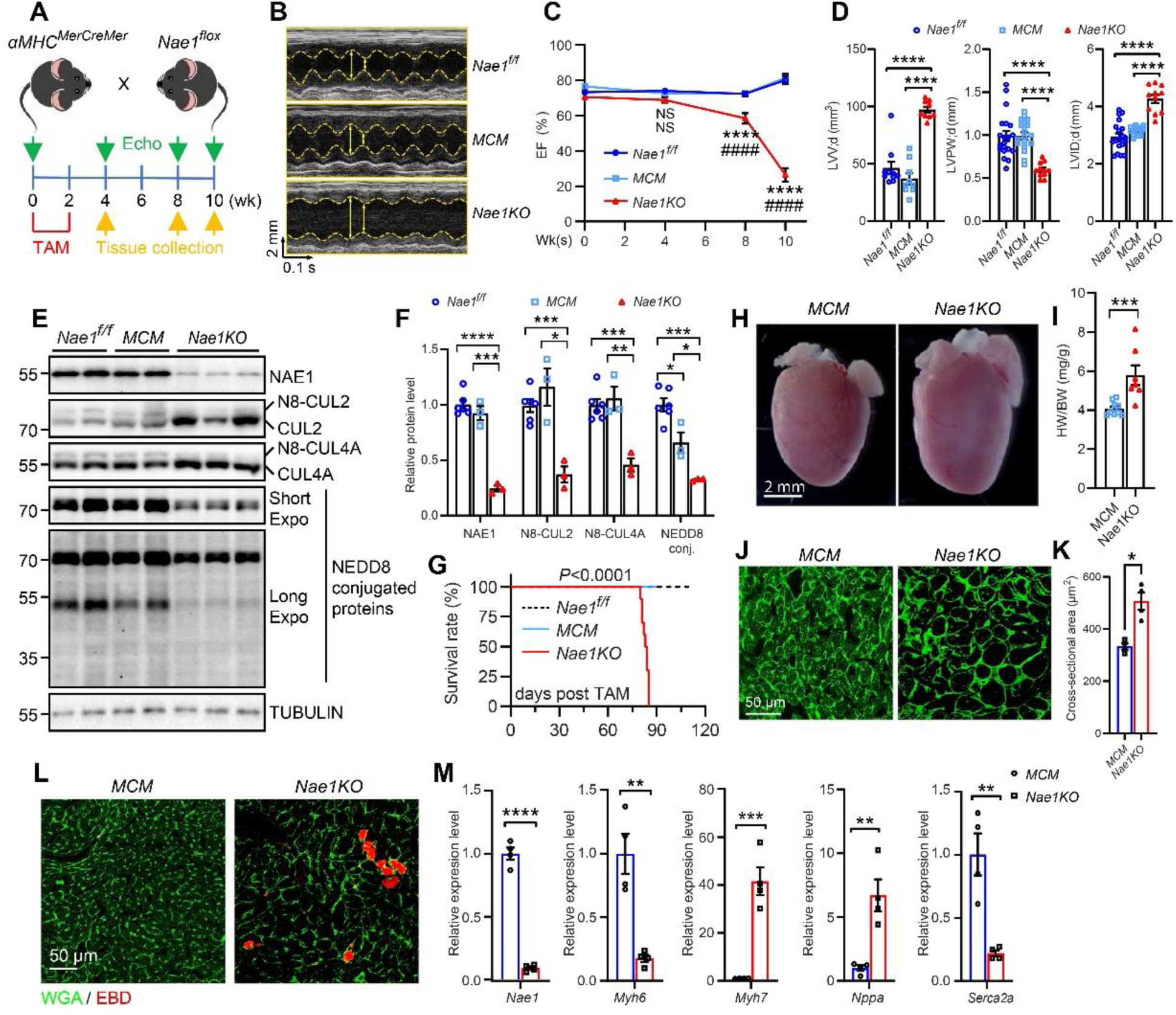
Neddylation is indispensable for post-mitotic murine hearts. **A**, scheme of the generation of *αMHC^MerCreMer^: Nae1^flox/flox^* inducible knockout mouse model and experiment timeline. TAM, tamoxifen; Echo, echocardiography. **B**, representative echocardiography images of indicated groups of mice at 10 weeks post TAM. **C**, temporal ejection fraction (EF) measured by echocardiography at indicated weeks (Wks). **D**, indicated echocardiography data measured at 10 weeks post TAM. LVV;d, left ventricle volume diastolic; LVPW;d, left ventricle posterior wall thickness diastolic; LVID;d, left ventricle internal diameter diastolic. **E,** western blot representing protein level as indicated, validating NAE1 knock out efficiency. N8, NEDD8; Expo, exposure; MCM, *αMHC^MerCreMer^*; KO, *αMHC^MerCreMer^*: *Nae1^flox/flox^*. Tissues collected at 4 weeks post TAM. **F**, quantification of western blots represented in **E** (NAE1^f/f^, n=6; MCM, n=3; KO, n=3). **G**, survial curve of indicated group of mice. **H**, gross morphology of MCM vs KO hearts collected at 8 weeks post TAM. **I**, heart weight/ body weight ratio (HW/BW) at collection (8 weeks post TAM). **J**, representative images of WGA staining of indicated group of mouse hearts collected at 10 weeks post TAM, scale bar is as indicated. **K**, quantification of cross-sectional area as shown in **J**. **L**, representative images of EBD staining and WGA staining of indicated group of mouse hearts collected at 10 weeks post TAM, scale bar is as indicated. **M,** qPCR analysis measuring relative mRNA level of indicated genes in MCM (n=4) vs KO (n=4). Tissues collected at 4 weeks post TAM. Statistical tests performed in panel **F, K** & **M** were nested t-test; **I** was student t-test; **C** & **D** were one-way ANOVA followed by post hoc Tukey’s multiple comparisons; **G** was log-rank test. *, 0.01<P< 0.05; **, 0.001<P< 0.01; ***, 0.0001<P< 0.001; ****, P< 0.0001; NS, not significant.

While NAE1KO hearts were initially functionally indistinguishable from NAE1^f/f^ and MCM hearts, they exhibited cardiac dysfunction at 8 weeks post TAM injections, with left ventricle (LV) ejection fraction dropped to ∼60% (Fig. 2C). These mice quickly progressed to severe dilated cardiomyopathy and heart failure by 10 weeks post-TAM, as indicated by significantly reduced ejection fraction (∼ 20%), decreased LV posterior wall thickness at diastolic state (LVPW; d), increased LV diastolic internal diameter (LVID; d), and increased left ventricle volume at diastolic state (LVV; d) compared to *Nae1^f/f^* and MCM mice (Fig. 2C-D). Eventually, all NAE1KO mice died within 90 days after TAM administration (Fig. 2G). Mutant hearts were significantly enlarged and has increased heart weight to body weight ratio(Fig. 2H-I). Histological analysis revealed increased cardiomyocyte cross sectional area (Fig. 2J-K). Evan blue dye (EBD) infiltration assay revealed increased EBD-positive cardiomyocytes, indicating necrotic cell death (Fig. 2L). qPCR analysis revealed significant downregulation of *Nae1* and *Myh6* and upregulation of *Myh7*, *Nppa*, and *Nppb* in mutant hearts (Fig. 2M). Thus, these data demonstrate an indispensable role of neddylation in post-mitotic murine hearts.

### Inhibition of neddylation exacerbates pressure overload-induced pathological cardiac remodeling and heart failure

Transverse aortic constriction (TAC) is a widely utilized surgery procedure to introduce pressure-overload mouse model to achieve cardiac hypertrophy and heart failure^33^. To determine the role of neddylation in regulation of pathological cardiac remodeling, we subjected CTL and NAE1KO mice to transverse aortic constriction (TAC) at 4 weeks after TAM administration- a time when their cardiac function is comparable (Fig. 3A). Echocardiography revealed that TAC surgery was successfully conducted (Fig. 3C & Sup. Fig. 3B), and TAC-induced pressure overload for 4 weeks led to an increase in LV wall thickness without altering LV internal diameter and ejection fraction in CTL mice, indicative of compensatory cardiac hypertrophy (Fig. 3B & Sup. Fig. 3A). In contrast, NAE1KO mice exhibited cardiac chamber dilation and had significantly reduced ejection fraction, suggesting accelerated cardiac decompensation and heart failure compared to CTL+TAC mice (Fig. 3B & Sup. Fig. 3A&C). Histological and immunohistological analyses showed significantly increased cardiomyocyte crosssection area, pronounced interstitial fibrosis and increased inflammatory cell infiltration in NAE1KO+TAC hearts (Fig. 3D-I). Taken together, these data suggested that inhibition of neddylation sensitizes the hearts to pressure overload-induced pathological cardiac remodeling and heart failure.

**Figure 3.**
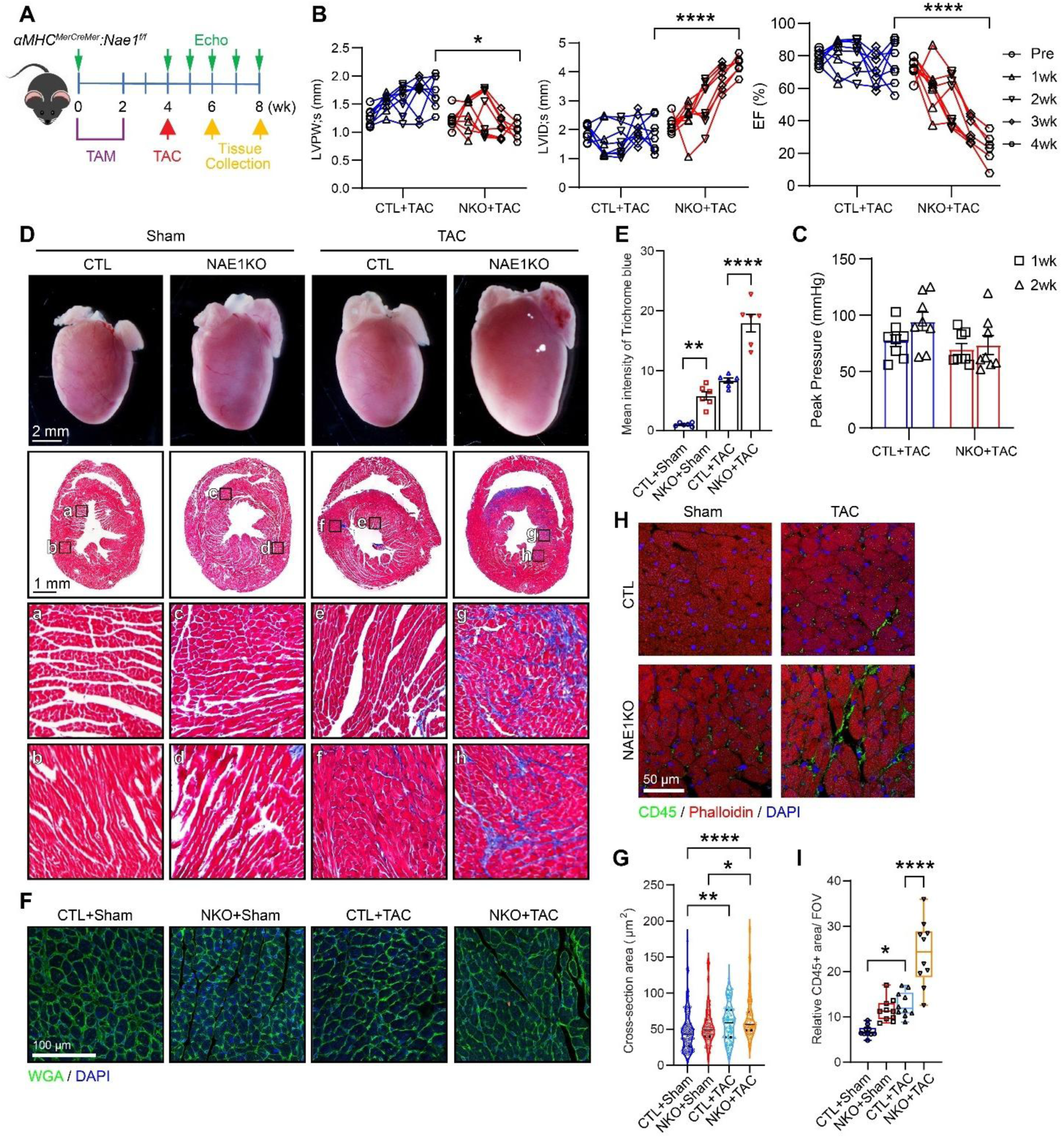
NAE1KO mice are sensitized to TAC-induced heart failure. **A**, Scheme of experiment design. **B**, Temporal change of LVPW; s, LVID; s, and EF before and after TAC (1, 2, 3, and 4 weeks as indicated). LVPW, left ventricle posterior wall thickness; LVID, left ventricle internal diameter; EF, ejection fraction; d, diastolic; s, systolic. **C**, Peak gradient pressure of TAC animals in 1 and 2 weeks after TAC. **D**, gross morphology (top row), trichrome staining (second row), and enlarged views (third and fourth rows) as labeled. **E**, quantification of fibrotic areas. **F**, cross-sectional wheat germ agglutinin (WGA) staining. **G**, quantification of cross-sectional areas. **H**, Representative confocal images of myocardial section immunostained with CD45 (green) and Phalloidin (red). **I**, Quantification of CD45 relative area per field of view (FOV). All scale bars represent length as indicated in each figure. Statistical tests performed in panels **B, E, G** and **I** were one-way ANOVA followed by post hoc Tukey’s multiple comparisons. *, 0.01<P< 0.05; **, 0.001<P< 0.01; ***, 0.0001<P< 0.001; ****, P< 0.0001; NS, not significant.

### RNA-seq analysis reveal dysregulated metabolic pathways in NAE1KO

To elucidate the underlying mechanism of NAE1KO-induced heart failure, we performed RNA-seq analysis on heart tissue of αMHC^MerCreMer^ (CTL, n=4) and αMHC^MerCreMer^: NAE1^flox/flox^ (NAE1KO, n=4) mice collected at 4 weeks after TAM injection, when cardiac functions showed no differences (Fig. 2C). Principal component analysis (PCA) showed two distinct groups of CTL and NAE1KO mice (Fig. 4A). In our RNA-seq analysis, there are 662 significantly downregulated genes and 849 upregulated genes (Fig. 4B, Fold Change >1.2, adjusted *P*<0.05). Gene set enrichment analysis (GSEA) revealed top enriched hallmarks, including upregulated sets such as reactive oxygen species (ROS) Pathway, myogenesis, inflammatory response, apoptosis, glycolysis, and downregulated sets such as oxidative phosphorylation, angiogenesis, adipogenesis, hypoxia, and fatty acid metabolism (Fig. 4C). Interestingly, glycolysis gene set was dysregulated in the NAE1KO group with an overall upregulated normalized enrichment score (NES) of 0.91 and nominal *P* value (Nom *P*) of 0.726 (Fig. 4D & G), whereas the fatty acid metabolism gene set (NES= -1.58, Nom *P*=0.003) and the oxidative phosphorylation gene set (NES=-0.99, Nom *P*=0.471) were drastically downregulated in NAE1KO (Fig. 4E, F, H & I). Consistently, qPCR analysis revealed that fatty acid metabolism genes *Acaa2, Acadvl, Dgat2, Ech1, Hadha, Mlycd, Ppara*, and *Eci1* were significantly decreased in NAE1KO mouse hearts (Fig. 4J). Together, these data suggest disrupted metabolic pathways in NAE1KO hearts.

**Figure 4.**
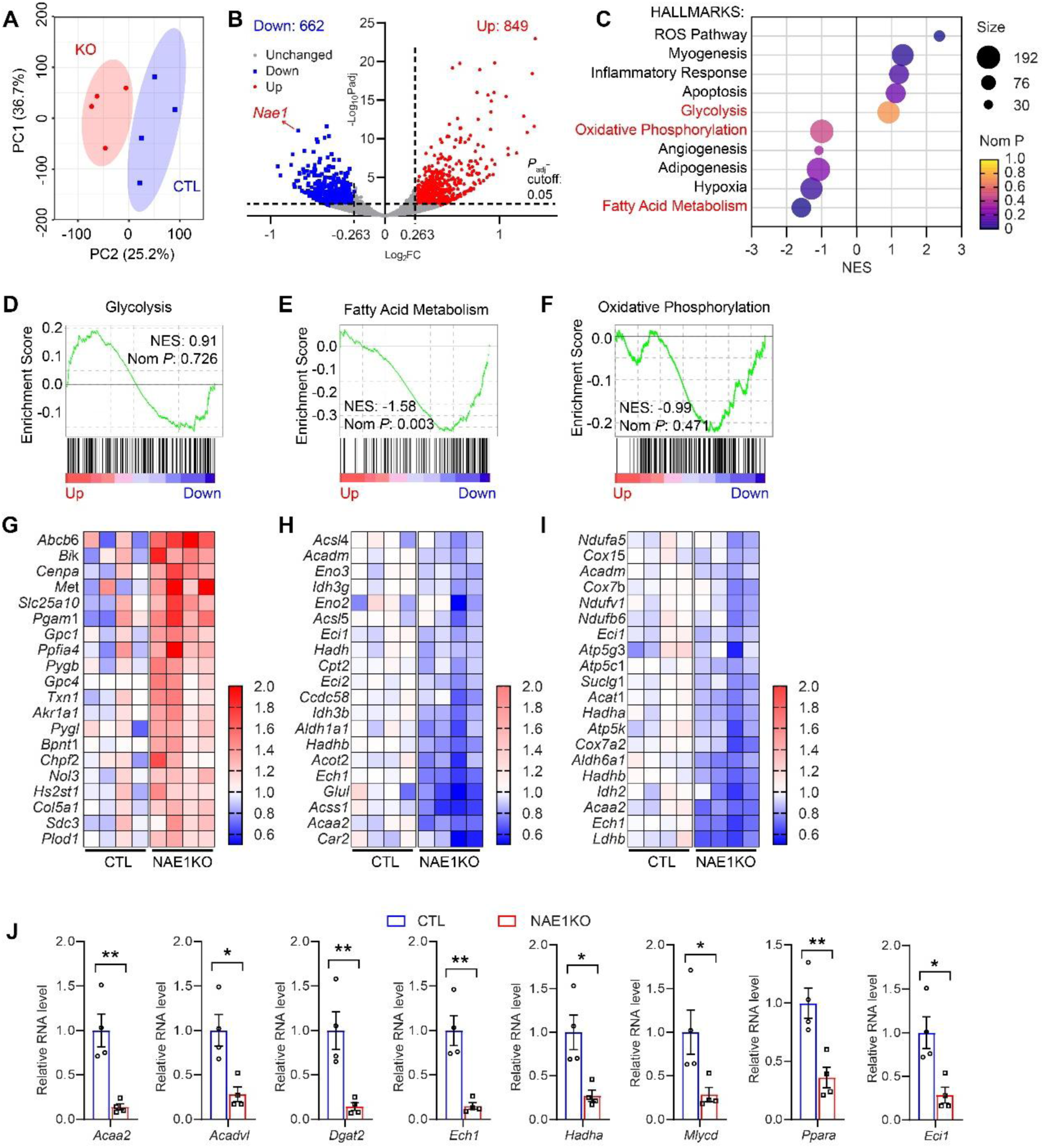
**RNA-seq analyses reveal metabolic and mitochondrial disruptions in the heart.** Ventricles of αMHC^MerCreMer^ mice as control (n=4, CTL) and αMHC^MerCreMer^: NAE1^flox/flox^ mice as NAE1KO (n=4, KO) were collected at 4 weeks post tamoxifen injection, and subjected to RNA-seq analyses. **A**, PCA analysis of 8 samples submitted. **B**, volcano plot showing upregulated (Up: 849 genes) and downregulated (Down: 662 genes) genes with thresholds of fold change (FC) more than 1.2 folds and adjusted P value smaller than 0.05. *Nae1* gene is as indicated in the plot. **C**, dramatically altered hallmark gene sets in gene set enrichment analysis (GSEA). Those gene sets labeled in red color were shown in following panels for more details. GSEA plots and heat maps of top 20 genes within that sets were shown as follow: glycolysis gene set (**D** & **G**), fatty acid metabolism gene set (**E** & **H**), and oxidative phosphorylation gene set (**F** & **I**). NES, normalized enrichment score; NomP, nominal P value. **J**, qPCR analyses of indicated genes in NAE1KO hearts to validate the RNA-seq analyses. Statistical tests performed in panel **J** were nested t-test. *, 0.01<P< 0.05; **, 0.001<P< 0.01; ***, 0.0001<P< 0.001; ****, P< 0.0001.

Other than the hallmarks pathway analyses, we also performed GSEA with mitochondrial gene sets (MitoCarta 3.0) (Sup Fig. 4A). It was shown that many mitochondrial pathways were downregulated in the NAE1KO group, including fatty acid oxidation (the top downregulated pathway, NES= -2.125, Nom P= 0.000, Sup Fig. 4B), lipid metabolism (NES= -1.528, Nom P= 0.005, Sup Fig. 4C)), TCA cycle (NES= -1.508, Nom P= 0.034, Sup Fig. 4D), and oxphos subunits pathway (NES= -1.417, Nom P= 0.022, Sup Fig. 4E). These data further imply the impaired mitochondrial genes and pathways in the NAE1KO hearts.

## Inhibition of neddylation impairs cardiac mitochondrial energetics and dynamics

Upon the RNA-seq analysis, it seems that inhibition of neddylation affected the metabolic pathways as well as mitochondrial genes (Fig. 4 & Sup. Fig. 4). To further investigate the role of neddylation in cardiac mitochondria, we first performed another GSEA on mitochondrial complexes. It was shown that genes within Complex I, III, IV, and V are downregulated and Complex II was upregulated in the NAE1KO hearts (Sup. Fig. 5A). Specifically, genes in Complex I are largely downregulated (NES= -0.917), such as NADH dehydrogenase family genes and NADH-ubiquinone oxidoreductase genes (Sup. Fig. 5B); genes in Complex IV also largely downregulated (NES= -1.221) especially the mitochondrial Cytochrome c oxidase family genes (Sup. Fig. 5D); while the genes in Complex II are altered bi-directionally (NES=0.641, Sup. Fig. 5C). The largely altered mitochondrial genes addressed our interest given the crucial interactions in between mitochondrial dysfunction and cardiomyopathies^5,6,11^.

**Figure 5.**
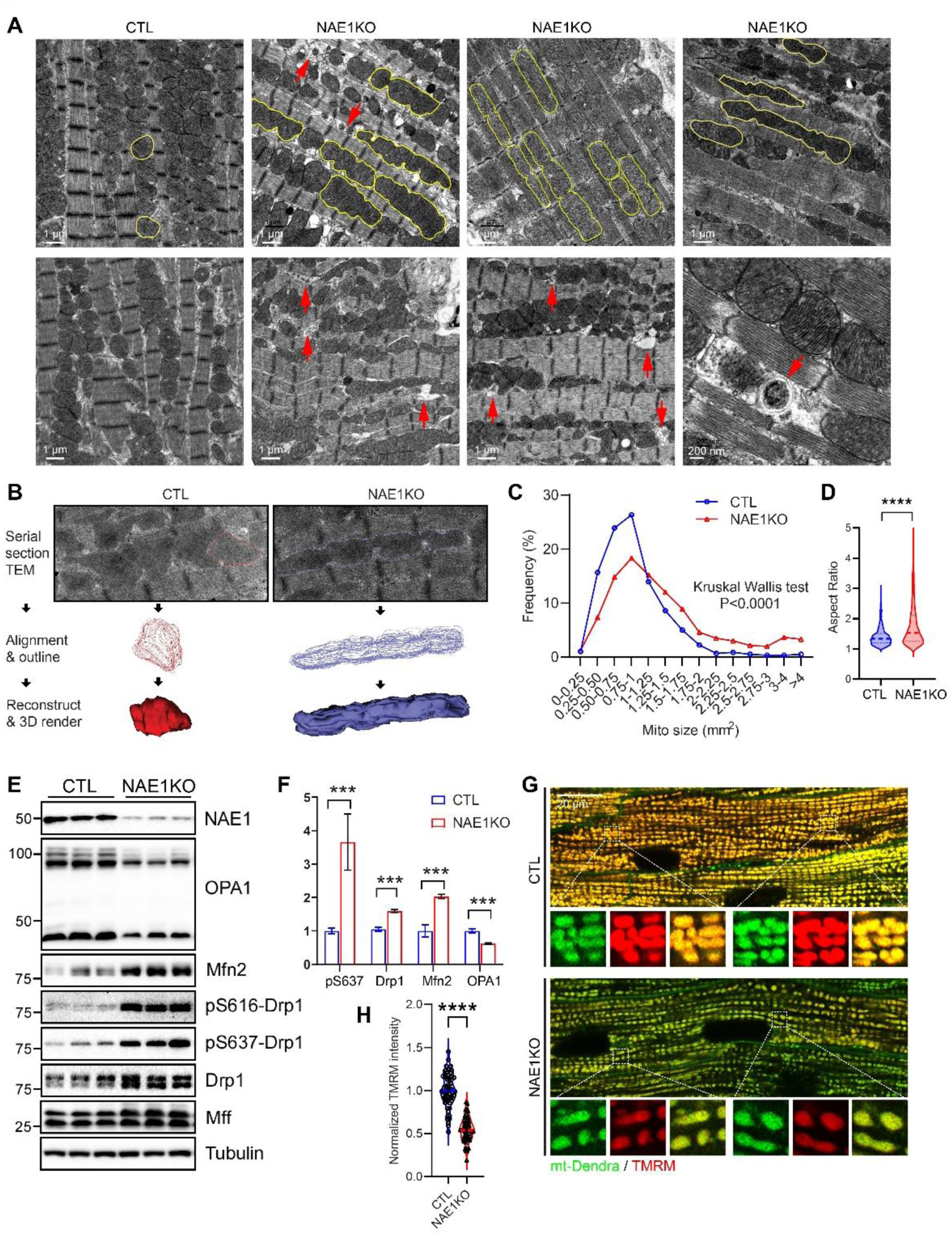
NAE1 inhibition impairs cardiac mitochondrial function and dynamics. **A**, transmission electron microscope (TEM) images of CTL and NAE1KO heart tissues. Yellow outline indicates mitochondria; solid red arrows indicate mitophagic vesicles. **B**, model of 3D-reconstruction from serial TEM images. **C**, quantification of frequencies of indicated mitochondrial size. **D**, aspect ratio of mitochondria in CTL and NAE1KO hearts. **E**, western blot (WB) of indicated proteins in CTL and KO hearts. **F**, quantification of WB shown in panel **E**. **G**, in situ confocal imaging showing TMRM staining in αMHCMerCreMer: COX8Tg/+ mice (CTL) and αMHCMerCreMer: NAE1flox/flox: COX8^Tg/+^ mice (NAE1KO). **H**, quantification of normalized TMRM intensity. All scale bars represent length as indicated in each figure. Statistical tests performed in panel **C** was Kruskal Wallis test, in panels **D** and **H** were student t test, panel **F** was one-way ANOVA followed by post hoc Tukey’s multiple comparisons. *, 0.01<P< 0.05; **, 0.001<P< 0.01; ***, 0.0001<P< 0.001; ****, P< 0.0001; NS, not significant.

To investigate the impact of NAE1KO on morphology of cardiac mitochondria, a careful transmission electron microscopy (TEM) analysis was performed to delineate the morphology of cardiac mitochondria in control and NAE1KO hearts. We performed 3Dreconstruction followed by serial scanning of TEM (Fig. 5A-B) and detected increased hyperfused mitochondria as well as mitophagic vesicles in the NAE1KO hearts compared to control hearts (Fig. 5A & Sup. Fig. 6A), indicated by increased frequency of larger sized mitochondria (Fig. 5C) and significantly increased aspect ratio (Fig. 5D). Consistent to the morphological observations, WB of NAE1KO heart tissue compared with control hearts reveals significantly dysregulated mitochondrial dynamic proteins in NAE1KO samples (Fig. 5E-F). Interestingly, elongated mitochondria were also observed in cardiac specific constitutive KO of Nae1 (NAE1cKO)^32^ P1 neonatal hearts (Sup. Fig. 6B). Meanwhile, the MLN-treated NRVCs with dramatic inhibition of neddylation (Sup. Fig. 6F) also displayed impaired mitochondrial dynamics protein levels (Sup. Fig. 6G-H). These data suggested that inhibition of neddylation induces hyperfusion of mitochondria in cardiomyocytes.

**Figure 6.**
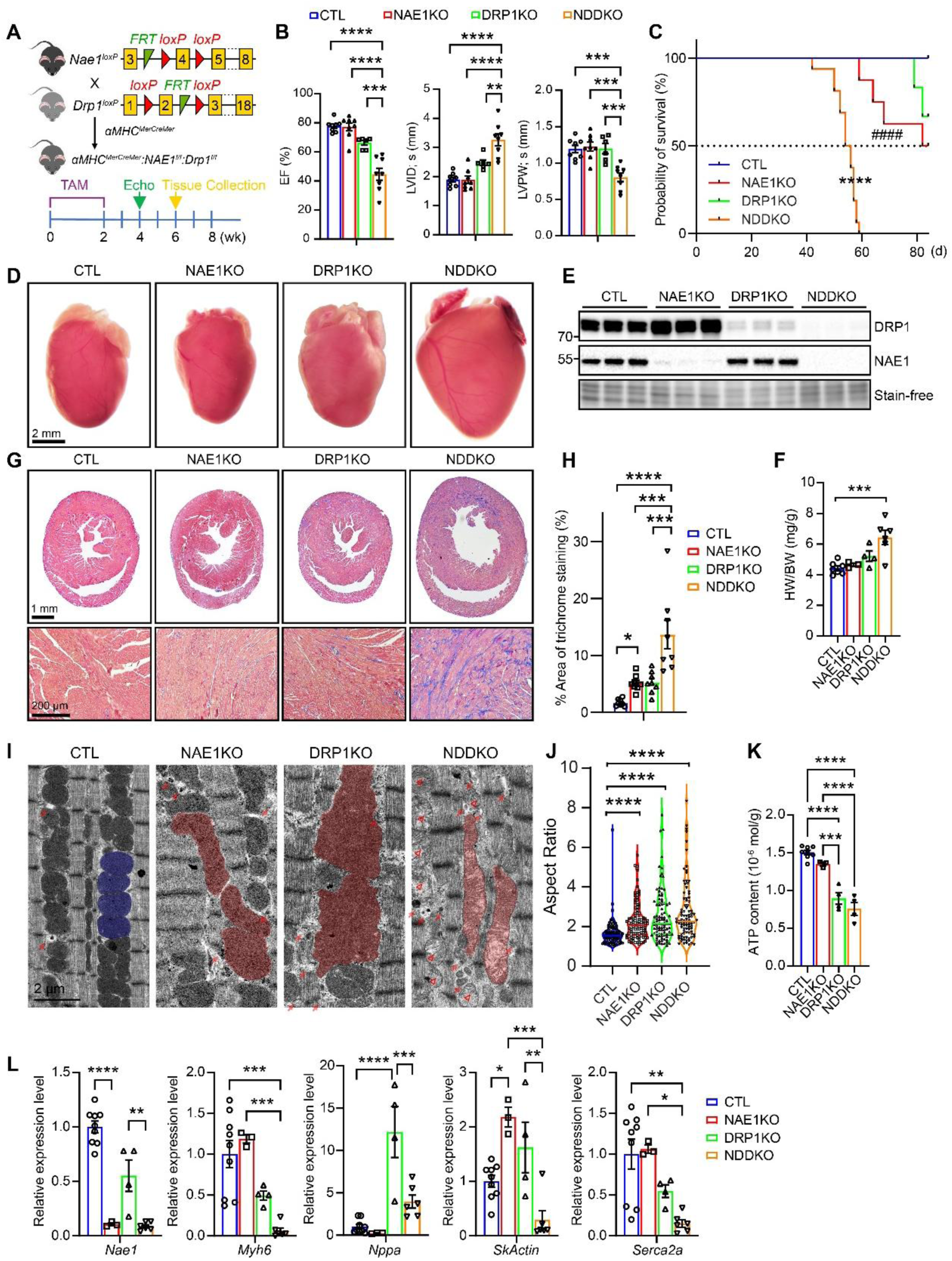
NAE1 deficiency exacerbates heart failure induced by DRP1 knockout. **A**, scheme of the generation of *αMHC^MerCreMer^:Nae1^f/f^:Drp1^f/f^* (*Nae1* and *Drp1* double knockout, NDDKO). **B**, Echocardiography measurements of indicated parameters at 4 weeks after TAM injection. EF, ejection fraction; LVAW, left ventricle anterior wall thickness; LVID, left ventricle internal diameter; LVPW, left ventricle posterior wall thickness; d, diastolic; s, systolic. **C**, survival curve of indicated mice. “*” represents comparison between CTL and NDDKO; “#” represents comparisons between NAE1KO and NDDKO. **D**, gross morphology of indicated mouse hearts. **E**, western blot of indicated proteins. **F**, heart weight/ body weight (HW/BW) ratio in indicated mice. **G**, trichrome staining (top row) and enlarged views (second and third rows) of yellow square fields. **H**, quantification of trichrome staining percentage in indicated groups shown in **F**. **I**, representative TEM images of indicated groups. Colored region indicates mitochondria. Solid red arrows indicate mitophagic vesicles; hollow red arrow heads indicate mitochondria with loose cristae. **J**, quantification of aspect ratio of indicated groups. **K**, ATP content measured in indicated group of heart tissues. **L**, relative mRNA level by qRT-PCR assay of indicated genes. All scale bars represent length as indicated in each figure. Statistical tests performed in panel **C** was log rank test, and all others were performed by one-way ANOVA followed by post hoc Tukey’s multiple comparisons. *, 0.01<P< 0.05; **, 0.001<P< 0.01; ***, 0.0001<P< 0.001; ****, P< 0.0001; NS, not significant.

The mitochondrial bioenergetics are also impaired in NAE1KO hearts. First, it was observed that the membrane potential indicated by TMRM staining was significantly reduced *in vivo* in NAE1KO hearts (Fig. 5G-H) . Meanwhile, the percentage of unhealthy isolated adult CMs (AdCMs) indicated by round-shape is increased in NAE1KO group (Sup. Fig. 7A-E) . In addition, the percentage of AdCMs completely losing mitochondrial membrane potential is also significantly increased in NAE1KO (Sup. Fig. 7C). Once treated with simulated ischemia-reperfusion (sI/R), the AdCMs isolated from NAE1KO hearts displayed even higher percentage of round-shape cells (Sup. Fig. 7E), indicating that inhibition of neddylation disrupts mitochondrial bioenergetics no matter under physiological condition or stressed condition in CMs. More direct evidence via Mito-Stress seahorse analysis within isolated AdCMs further suggested a crucial role of neddylation in cardiac mitochondria (Sup. Fig. 7I). It was observed that the basal respiration, maximum respiration, proton leak, ATP production, and spare respiration capacity are all significantly decreased in MLN-treated AdCMs (Sup. Fig. 7J). Notably, the activity of mitochondrial Complex I is also repressed in NAE1KO hearts (Sup. Fig. 7K). These data together suggested a dispensable role of neddylation in cardiac mitochondrial respiration and bioenergetics.

**Figure 7.**
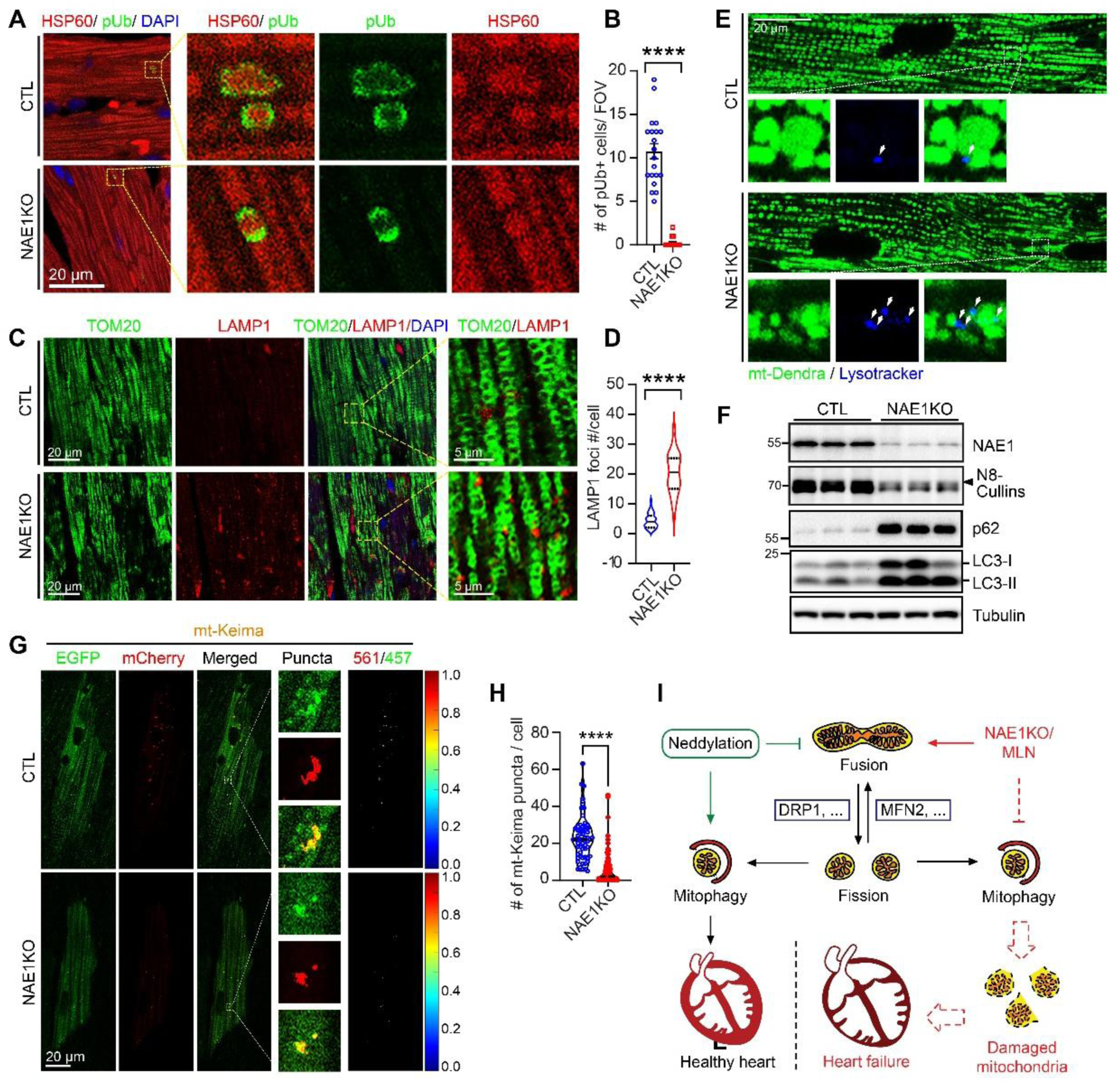
Neddylation is required by mitochondria turnover. **A**, Immunofluorescent staining of longitudinal sections in hearts from *αMHC^MerCreMer^* mice (CTL) and *αMHC^MerCreMer^: Nae1^flox/flox^* mice (NAE1KO) against HSP60, phosphorubiquitin (pUb) at Ser-65. **B**, quantification of number of cells with pUb positive staining per cell as represented in **A**. **C**, Immunofluorescent staining of longitudinal sections in hearts from CTL and NAE1KO against TOMM20 and LAMP1. **D**, quantification of number of LAMP1 foci per cell as represented in **C**. **E**, representative images of in situ confocal scanning of *Cox8^mtDendra^* mice stained with LysoTracker. **F**, western blot representing protein levels as indicated. N8-Cullins represent neddylated Cullins band. **G**, hearts from indicated mice were infected with AAV-Mito-Keima and subjected to in situ confocal microscopy. The ratio map was calculated by ratio of intensities within indicated channels and rendered in MATLAB as heatmap. **H**, quantification of number of positive puncta per cell as represented in **G**. **I**, a proposed model how neddylation impairs adult heart via regulating mitochondria dynamics and turnover. Scale bars are as indicated in each specific panel. Statistical tests performed in panels **B, D** and **H** were nested t test. *, 0.01<P< 0.05; **, 0.001<P< 0.01; ***, 0.0001<P< 0.001; ****, P< 0.0001; ns, not significant.

### Depletion of NAE1 exacerbates DRP1KO-induced cardiomyopathy

As observed in NAE1KO hearts, interestingly, even though the mitofusion protein MFN2 is increased supporting the observation of elongated mitochondria, another mitofusion regulator OPA1 is decreased for both long form and short form in the heart but not in NRVCs, while DRP1 total protein level, together with its activating and inhibiting phosphorylated forms pS616-DRP1^34,35^ and pS637-DRP1^36^ levels are consistently increased both *in vivo* and *in vitro* (Fig. 5E-F & Sup. Fig. 7G-H). These data implied that neddylation potentially regulates mitochondrial dynamics via interplaying with mito-dynamic proteins, such as DRP1 and MFN2.

To investigate the interplays between neddylation and DRP1, we generated an inducible double knockout mouse model that depletes both NAE1 and DRP1 specifically in cardiomyocytes (Fig. 6A). Similar to previous findings^8^, αMHC^Mer-Cre-Mer^ (MCM) induced cardiac specific DRP1 knockout mouse (DRP1KO) gradually exhibits reduced ejection fraction after TAM injection (Sup. Fig. 8A). At 4 weeks post TAM injection, echocardiography showed that only the MCM-NAE1-DRP1 double knockouts (NDDKO) have significantly decreased ejection fraction (∼40% vs ∼70-80% of CTL, NAE1KO, and DRP1KO), increased internal diameters, and decreased wall thicknesses (Fig. 6B & Sup. Fig. 8C). Survival analysis revealed that NDDKO mice reached lethality significantly faster than all other groups, and none of them could survive until 60 days after TAM injection (Fig. 6C). In addition, during collection at 6 weeks after TAM injection, only the NDDKO showed significantly enlarged heart size, but not NAE1KO or DRP1KO (Fig. 6D & F). These data suggest that depletion of NAE1 largely escalates DRP1KO-induced heart failure and early mortality.

To further investigate the mechanism of aggravated cardiomyopathy, we first validated the knockout efficiency of both NAE1 and DRP1 (Fig. 6E). Next, we observed significantly increased fibrosis of the NDDKO heart by trichrome staining (Fig. 6G-H), which is consistent with dysregulated fetal genes expression (Fig. 6L) and fibrotic genes expression (Sup. Fig. 8D). This suggests that the NDDKO hearts went through severe cardiac remodeling during the disease progression. Furthermore, we investigated mitochondrial morphology within CMs via TEM and observed accumulated hyperfused mitochondria (Fig. 6I & Sup. Fig. 9A) indicated by significantly increased aspect ratio, form factor, and decreased circularity (Fig. 6J, Sup. Fig. 9B-C, respectively) of mitochondria. Intriguingly, despite that the NDDKO has a trend of more severe hyperfusion, it did not exceed NAE1KO or DRP1KO significantly, while producing significantly less ATP content (Fig. 6K). These data suggest that disrupted mitochondrial dynamics caused by NAE1KO may not solely contribute to the exacerbated cardiomyopathies.

### Double knockout of NAE1 and MFN2 leads to cardiomyopathy and lethality

As we identified accumulated MFN2 in NAE1KO hearts (Fig. 5F), together with the hyperfused mitochondria in CMs of NAE1KO (Fig. 5A-D), we next tested whether double knockout of NAE1 and MFN2 in the heart could rescue the mitochondrial dysfunction and heart failure by generating an αMHC^Mer-Cre-Mer^-induced cardiac specific NAE1 and MFN2 double knockout mouse model (NMDKO, Sup. Fig. 10A). The CM-restricted knockout appeared to effectively deplete NAE1 and MFN2 proteins (Sup. Fig. 10B). Even though the NMDKO eventually exhibit similarly reduced ejection fraction, interventricular septum, and enlarged internal diameters as NAE1KO at late stage of disease progression (8-10 weeks), it did perform a protective role at early stage (6 week) (Sup. Fig. 10C-E). Subsequently, the NMDKO mice showed significantly earlier lethality compared to NAE1KO (Sup. Fig. 10F). During tissue collection at 10 weeks post TAM injection, enlarged heart size and increased lung weight similar to NAE1KO were also observed in NMDKO mice (Sup. Fig. 10G & H). Taken together, these data implied a potential early-stage protection by inhibiting mitochondria hyperfusion in NAE1KO hearts, while substantiated that NAE1KO-induced cardiomyopathy may not be exclusively attributed to mitochondrial dynamics dysregulation.

### Neddylation is required by mitochondria turnover

Other than hyperfused mitochondria, we also observed significantly increased mitophagic vesicles within the NAE1KO hearts in TEM (Fig. 5A, Fig. 6I & Sup. Fig. 6A & 9A). Notably, only NAE1KO and NDDKO leads to increased mitophagic vesicles, but not DRP1KO (Sup. Fig. 9D). Thus, we next hypothesize that neddylation is crucial for mitophagy.

pS65-Ubiquitin (pUb) is previously reported to modify outer mitochondrial membrane (OMM) proteins upon mitophagy^37^. It was observed within NAE1KO heart tissues that the pUb colocalizing with mitochondrial marker HSP60 is significantly decreased compared to control (Fig. 7A & B), which suggests a decrease in mitophagy. On the other hand, lysosomal membrane protein LAMP1 significantly increased in NAE1KO (Fig. 7C & D), suggesting an increased number of lysosomes at whole-cell level. This is consistent with the in vivo live tissue scanning that lysosome number indicated by Lysotracker staining also increased in NAE1KO heart (Fig. 7E), and the live cell imaging of isolated AdCMs from NAE1KO hearts that intermyofibrillar lysosome also significantly increased (Sup. Fig. 11A-B). In addition, WB analysis demonstrated increased whole cell levels of lysosomal proteins p62 and LC3-II in NAE1KO hearts (Fig. 7F), while treatment of MLN in NRVCs dramatically decreased CCCP-induced pUb (Sup. Fig. 11E&H) as well as mitochondrial p62 and pUb (Sup. Fig. 11H&J). Meanwhile, mitophagy-flux assay also revealed decreased mitochondrial LC3-II level upon MLN treatment (Sup. Fig. 11K). Consequently, MLN-treated NRVCs displayed increased mitochondria content upon mitochondrial stress (Sup. Fig. 11F). These data suggest that given the lysosome level are upregulated in NAE1KO, the phosphorylation of mitochondrial Ub is repressed upon inhibition of neddylation.

To further investigate whether the increased number of mitophagic vesicles observed in NAE1KO hearts perform well-maintained functions, we introduced AAV9-cTnT-mtKeima as a mitophagic function readout. Effective mitophagy requires the microenvironment within mitophagic vesicle to be acidic, which would turn mt-Keima from green to red fluorescence and thus mitophagy could be represented by high red/green ratio of mt-Keima^38,39^. It was observed that the puncta number of mt-Keima with high red/green ratio in NAE1KO are significantly decreased in live tissue scanning *in vivo* (Fig. 7G-H). Consistently, MLN treatment in CCCP-stress-induced NRVCs significantly decreased Ad-mt-Keima red/green ratio (Sup. Fig. 11C-D). We also utilized another mitophagy marker Cox8^EGFP-mChery^^40^ in H9C2 cells and observed similar readout (Sup. Fig. 11G). These data suggest that neddylation is critical for effective mitophagy, especially under stressed conditions.

To carefully delineate the impact of different conditions of MLN treatment, a time course CCCP analysis was performed and we uncovered that under long term treatments of CCCP (12h and 24h), MLN treatment showed better effect on inhibiting mitophagy, indicated by largely decreased pUb in mitochondrial fraction (Sup. Fig. 11J). In addition, a dose-dependent experiment in NRVCs revealed that MLN treatment can effectively inhibit CCCP-induced mitophagy at as low as 0.1 µM (Sup. Fig. 12A).Furthermore, to test the impact of inhibition of neddylation for mitophagy in different kinds of mitochondrial stress, we applied CCCP treatment, starvation, and hypoxia stresses to NRVCs, and observed that only CCCP-induced mitophagy, indicated by pUb level, could be inhibited by MLN-treatment (Sup. Fig. 12B). These data validated the indispensable role of neddylation in mitophagy under CCCP-induced stress conditions.

Cullins (CULs) are the most well-established substrates of neddylation, participating in many cellular processes such as proliferation, programmed cell death, metabolism, etc.^23,24,41–43^. It was observed that several CULs are localized to mitochondria, including CUL2, CUL3, CUL4a and CUL4b (Sup. Fig. 11I). To investigate the mechanism underlying the role of neddylation in mitophagy, we next screened multiple CULs with siRNA treated NRVCs under CCCP treatment. It was shown that knocking down CUL2 and CUL4b with two different siRNAs can constantly decrease CCCP-induced upregulation of pUb, while CUL1 and CUL9 showed some inconsistence (Sup. Fig. 13A), suggesting that CUL2 and CUL4b might be the downstream target of neddylation in regulating mitophagy. Seahorse analysis showed that unlike treatment of MLN that inhibits all neddylations by inactivating the E1 enzyme, siCUL2 or siCUL4b, as a target of neddylation, did not significantly impair the respiration rate of mitochondria in NRVCs (Sup. Fig. 13D-E). However, siRNA of CUL2 and CUL4b could both inhibit mitochondrial targeted pUb under both regular condition and CCCP-induced stress conditions (Sup. Fig. 11B-C). These data suggest that CUL2 and CUL4b might be the downstream target of neddylation in regulating mitophagy, but not mitochondrial bioenergetics.

Taken all results together, we propose that neddylation plays crucial role in fine-tuning mitochondria fusion/ fission dynamics and mitophagic turnover in healthy hearts; upon inhibition, impaired neddylation leads to hyperfused mitochondria and defected mitophagy, which in turn leads to accumulated damaged mitochondria in the heart and subsequent cardiomyopathies like heart failure (Fig. 7I).

## DISCUSSION

In this study, we elucidated the pivotal role of neddylation in maintaining cardiac mitochondrial homeostasis and function under both physiological and pathological conditions. Neddylation appears imperative for moderating mitochondrial dynamics and turnover, as demonstrated by its impact on mitochondrial dynamic modulators like Drp1 and Mfn2, and in preserving mitophagy integrity. Disruption in neddylation perturbs mitochondrial morphology, leading to hyperfusion and mitochondrial damage, alongside compromised mitochondrial turnover. These mitochondrial dysfunctions result in reduced energy production, heart failure, and ultimately, lethality. Our findings unveil a previously overlooked significance of neddylation in cardiac health in post-mitotic hearts, offering novel perspectives on the pathogenesis of cardiomyopathies.

Neddylation was previously reported to mainly contribute to regulation of cell proliferation and tumorigenesis, and thus considered as an antitumor target therapeutically^44,45^. The small molecule inhibitor of NAE, MLN4924 (Pevonedistat), is found to exhibit potent anticancer activity, and various clinical trials targeting multiple malignancies were undergoing^46,47^. Thus, certain concerns were raised regarding the off-target effects of MLN4924 on normal cells, including CMs in the developing heart. We have previously reported that neddylation contributes to cardiac development and maturation by promoting CM proliferation via Hippo-YAP signaling^25^ as well as mediating cardiac metabolic maturation via HIF1α signaling pathway^27^. In addition, we revealed that transient treatment of MLN4924 during neonatal stage predisposed the heart to stress sensitization in rats, which addressed the concerns of MLN4924 treatment in children and pregnant women^26^. Here in this paper, we raised another concern regarding cardiotoxicity of therapeutic strategies targeting neddylation pathway even in postmitotic hearts, as inhibition of neddylation disrupts cardiac mitochondrial dynamics and turnover, which are crucial for adult CMs.

Mitochondria play a central role in the heart and contribute to ∼30% of adult heart content^48,49^. To maintain normal function and high energetic efficiency, mitochondria in CMs are highly dynamic organelles whose structure and function are regulated by biogenesis, fusion, fission, and mitophagy^6,7^, coordinated by key regulators such as Drp1 for fission^8^ and Mfn1/2 for fusion^9,10^. Drp1, responsible for mitochondrial fission, maintains energy balance and mitochondrial quality in high-demand cardiac cells. Mutations like A395D in humans^50,51^ and C452F in Python mice^52,53^ disrupt Drp1 GTPase function, impair self-assembly, and lead to severe metabolic defects and cardiomyopathy, highlighting the essential role of Drp1 in cardiomyopathies. However, the role of posttranslational modification in Drp1-linked cardiomyopathies remains largely unknown. Interestingly, we found that inhibition of neddylation exacerbates heart failure caused by Drp1CKO (Fig. 6), implying a crucial role of neddylation in Drp1-linked cardiomyopathies. Likewise, Mfn2 regulates mitochondrial fusion and selective mitophagy, and the R400Q mutation specifically impairs mitophagy, resulting in mitophagic cardiomyopathy due to the accumulation of dysfunctional mitochondria^54^. Previous report suggested that cardiac restricted KO of Mfn2 results in enlarged and morphologically abnormal mitochondria, which exhibit increased resistance to calcium-induced mitochondrial permeability transition pore (MPTP) opening, while it does not lead to severe heart failure^55^. Interestingly, we found that inhibition of neddylation drastically disposed Mfn2CKO mice into severe heart failure (Sup. Fig. 10), suggesting an indispensable role of neddylation within Mfn2-related cardiomyopathies. Taken together, our findings support an elevated concern of the cardiotoxicity on inhibition of neddylation, especially within those related to metabolic disorders.

While inhibition of neddylation leads to disrupted mitochondrial dynamics and turnover, which subsequently leads to heart failure, we do not exclude the potential involvement of other important pathways. It was suggested that various other cellular pathways were found disturbed in NAE1KO heart, including upregulated ROS pathway, myogenesis, inflammatory response, apoptosis, and downregulated angiogenesis and adipogenesis pathways (Fig. 4C). Disrupted ROS pathway, as well as its interplay with mitochondria dysfunction, has been largely implicated in cardiomyopathies^56^. More importantly, perturbation of inflammatory responses may precipitate cardiomyopathies, including dilated cardiomyopathy and myocarditis, culminating in myocardial injury and compromised cardiac function^57^. In this study, we focused on the impact of neddylation on mitochondria in the heart, while raised intriguing questions regarding other cellular pathways that might be involved in neddylation mediated cardiac maintenance for future directions.

In a canonical model of removing damaged mitochondria via mitophagy involving PINK1/PARKIN pathway, mitophagy is initiated when mitochondrial membrane potential is lost, stabilizing PINK1 on the outer mitochondrial membrane, where it phosphorylates ubiquitin (pS65-Ub) and activates PARKIN, an E3 ubiquitin ligase^58^. PARKIN amplifies ubiquitin chain formation on mitochondrial proteins, facilitating recruitment of autophagy receptors like p62^59^. p62 bridges ubiquitinated mitochondria to autophagosomal LC3-II through its LIR domain^60,61^, enabling sequestration of damaged mitochondria into autophagosomes. Subsequently, autophagosomes fuse with LAMP1-positive lysosomes to form autolysosomes^62^, where hydrolytic enzymes degrade the mitochondria, recycling their components to maintain cellular homeostasis. In this study, we observed disrupted mitochondria function and dynamics upon inhibition of neddylation (Fig. 5), which potentially contributed to upregulated damaged mitochondria. Under normal conditions, damaged mitochondria with membrane potential loss triggers mitophagy and removal of damaged mitochondria. However, inhibition of neddylation causes failed initiation of Ub phosphorylation on mitochondria (Fig. 7A, Sup. Fig. 11), leading to accumulated damaged mitochondria and subsequent stress in CMs (Fig. 5A, Sup. Fig. 6), which likely induces compensatory accumulation of lysosomes indicated by upregulated LAMP1 (Fig. 7C) and formation of autophagic vesicles indicated by upregulated p62 and LC3II protein level (Fig. 7F). This proposed model above explains the significantly downregulated mitophagy in CMs with neddylation inhibition, indicated by decreased mt-Keima as a direct mitophagy marker (Fig. 7G).

On the other hand, we do not exclude the possibility that neddylation may regulate mitophagy in multiple models other than proposed in this study. It is reported previously that SCF^FBXL^^4^, an SKP1/CUL1/F-box ubiquitin ligase complex regulated by neddylation, localizes to the mitochondrial outer membrane in unstressed cells, where it constitutively ubiquitinates and degrades mitophagy receptors NIX and BNIP3 to suppress basal mitophagy^63^. In addition, a back-to-back study of the previous one identified the role of VHL, a cullin-RING ligase substrate receptor regulated by neddylation, in mitophagy^64^. This may explain the upregulated mitophagy we observed upon neddylation inhibition in CMs under basal condition (Sup. Fig. 12A).

In summary, our study revealed a crucial role of neddylation in finetuning mitochondrial dynamics and turnover in CMs, safeguarding cardiac homeostasis under both physiological and pharmacological conditions, which provides important insights in potential clinical impact on cardiomyopathies and addressed attention to the care of cardiotoxicity of certain chemotherapy. Future exploration in detailed mechanisms of our proposed model may facilitate the discovery of specific therapeutic targets for cardiomyopathies.

## METHODS and MATERIALS

### Animals

All animal experiments were approved by the Augusta University Institutional Animal Care and Use Committee. A transgenic mouse line bearing a cardiac specific knockout of NAE1 (NAE1cKO) was described previously^25^. A “knockout-first” allele^65^ (NAE1^neoflox^) was rederived from targeted frozen mouse embryos (EPD0441_1_C08) created by Eucomm. The mutant mice were first bred to ACTB^FLP^ transgenic mice^66^ (the Jackson Laboratory, #005703) to remove Frt-flanked neo cassette, then to *αMHC^MerCreMer^* mice^31^ (the Jackson Laboratory, #005657) to generate tamoxifen-inducible cardiomyocyte-restricted NAE1 knockout (NAE1KO) mice. *Drp1^flox^* mice were provided by Drs. Tohru Fukai and Masuko Ushio-Fukai, Augsuta University. *Mfn2^flox^* mice were obtained from JAX (B6.129(Cg)-Mfn2tm3Dcc/J, # 026525). These mice were maintained in the C57BL/6J inbred background for our studies. Male and female mice were equally included in all experiments. No significant sex differences were observed.

### Echocardiography

Mice at the age of 8-12 weeks were rendered unconscious through the inhalation of isoflurane gas, using a concentration of 2.5% for the initial sedation and 1.5% for ongoing sedation, administered through a nose cone. The depth of anesthesia was assessed by performing a toe pinch test. Heart imaging and motion capture were carried out using a VEVO 2100 echocardiography system equipped with a 30MHz transducer, produced by Visual Sonics. Analysis of the left ventricle’s shape and functionality was conducted later using VEVO 2100’s dedicated software. For the echocardiography of awake young mice, they were carefully held in place on the examination platform using adhesive tapes as previously described^25,27^.

### RNA-seq analysis

Mouse ventricles of 4 CTL mice and 4 NAE1KO mice were collected at age as described and stored following RNAlater (Invitrogen) manufacture protocol at -80 °C. The mRNA of the tissue was isolated through TRIzol-based protocol (Invitrogen). Isolated RNA was subjected to RNA quality control and RNA-sequencing analysis performed by Genome Technology Access Center (GTAC) via Next Generation Sequencing (NGS) as previously described^27^.

### Histology and immunohistochemistry analysis

For histological analysis, paraffin-embedded tissues were sectioned at 5 µm and processed for H&E or Trichrome staining. For immunofluorescence, cryosections underwent antigen retrieval in preheated sodium citrate buffer (pH 6.0, 98 °C) for 10 min using the PT Link system (Dako). In the case of cryo-sectioned tissues, deparaffinization and antigen retrieval were replaced by incubation in 1% Triton X-100 in PBS for 10 min at room temperature. Following blocking with 10% non-immune goat serum (Thermo Fisher Scientific) to reduce non-specific binding, sections were incubated with primary antibodies overnight at 4 °C and then with the appropriate Alexa-Fluor-conjugated secondary antibodies (Thermo Fisher Scientific) for 1 h at room temperature. Nuclei were counterstained with DAPI (Sigma), and slides were mounted using VECTASHIELD antifade mounting medium (Vector Laboratories). Images were acquired using an Olympus BX41 microscope (Olympus) or a Zeiss Upright 780 confocal microscope (Zeiss).

### Isolation and culture of adult cardiomyocytes

Adult ventricular cardiomyocytes were isolated from 8-12-week-old NAE1KO or control mice using a modified protocol previously reported^67^. Mice were anesthetized with inhaled isoflurane, and hearts were rapidly excised and transferred to ice-cold, calcium-free Tyrode’s solution to halt metabolic activity. The aorta was cannulated, and hearts were mounted on a Langendorff perfusion system. Retrograde perfusion was carried out at 3 mL/min and maintained at 37 °C. Hearts were first perfused with calcium-free Tyrode’s solution for 5 min to remove blood and equilibrate the tissue. Enzymatic digestion was then initiated by perfusion with Tyrode’s solution containing 0.5 mg/mL collagenase type II and 0.02 mg/mL protease for 15 min. Following digestion, ventricles were dissected, gently minced in a stop solution (Tyrode’s solution supplemented with 0.1 mM CaCl₂ and 1% BSA), and the resulting suspension was filtered through a 100µm nylon mesh. Calcium was gradually reintroduced by stepwise additions of CaCl₂ from 0.1 mM to 1.8 mM over 30 min. Isolated cardiomyocytes were resuspended in M199 medium containing 0.1% BSA, 10 mM 2,3-butanedione monoxime (BDM), 1× ITS, 1× CD lipids, 100 U/mL penicillin, and 100 μg/mL streptomycin, and plated onto dishes pre-coated with 10 μg/mL laminin. Cultures were maintained at 37 °C in 5% CO₂. Cell viability was assessed by trypan blue exclusion, and cell morphology was verified by phase-contrast microscopy. Only rod-shaped cardiomyocytes with intact striations were used for downstream experiments.

### In situ cardiomyocyte scanning

Hearts from NAE1KO and CTL mice were isolated and either perfused with LysoTracker or left untreated, then subjected to in situ confocal microscopy as previously described previously^68^. Briefly, intact mouse hearts were Langendorff-perfused at room temperature using Tyrode’s solution composed of 137 mM NaCl, 5.4 mM KCl, 10 mM HEPES, 10 mM glucose, 1 mM MgCl₂, and 0.33 mM NaH₂PO₄, with the pH adjusted to 7.4 using NaOH and oxygenated with 95% O₂ and 5% CO₂ during experiments. The perfusion solution included 10 μM LysoTracker (Thermo Fisher), a fluorescent dye that selectively stains acidic organelles like lysosomes, and was applied for 20 minutes at 37°C. The fluorescent signals from epicardial myocytes were analyzed in situ using a confocal microscope (STELLARIS 8, Leica Microsystems).

### Transmission electron microscopy and 3D re-construction

Transmission electron microscopy (TEM) sample preparation and imaging were performed at the Electron Microscopy and Histology Core at Augusta University. Briefly, left ventricular tissue was fixed at room temperature for 24 h in 3.5% glutaraldehyde in Sorensen’s phosphate buffer. Following fixation, samples were dehydrated, and ultrathin sections were prepared using a Leica EM UC6 ultramicrotome and mounted onto 200-mesh copper grids. Sections were stained with 2% aqueous uranyl acetate for 20 min, followed by lead citrate staining. Imaging was conducted on a JEOL JEM1400 Flash electron microscope, and representative images were selected to illustrate key ultrastructural features. Approximately 200 consecutive sections were acquired for 3D reconstruction. Images were aligned and reconstructed using a previously published platform (Reconstruct)^69^ and subsequently edited for presentation in Adobe Illustrator.

### Culture of neonatal cardiomyocytes and cell lines

Neonatal rat ventricular cardiomyocytes (NRVCs) were isolated using the Neonatal Cardiomyocyte Isolation System (Worthington) according to the manufacturer’s instruction^70^. Hypoxic conditions were established by placing cells in a humidified N₂ chamber supplemented with 5% CO₂ and 1% O₂ at 37 °C for 24 h. HEK293 cells were maintained in DMEM supplemented with 10% fetal bovine serum at 37 °C in 5% CO₂. The pShuttle-CMV-SF-NEDD8 plasmid (encoding Strep- and FLAG-tagged NEDD8) was generated as previously described^70^ and transfected into cells using X-tremeGene HP DNA transfection reagent (Roche) following the manufacturer’s protocol. Cells were harvested 48-72 h post-transfection or as otherwise indicated. Where applicable, cells were treated with vehicle controls (DMSO or bovine serum albumin), 1 µM MLN4924 (Active Biochem), 100 nM bortezomib (Enzolife), or 1 nM Echinomycin (Thermo Fisher).

### Protein extraction and western blot analysis

Protein was extracted from ventricular myocardium tissues or cultured cells, concentration determined with BCA reagents (Thermo Fisher Scientific), and SDSPAGE, immunoblotting, and densitometry analysis were performed as previously described^71^. Primary antibodies used in this study are listed in **Supplemental Table 1**.

### Triglyceride (TG) measurement

Cells were lysed in 1% Triton X-100 in PBS. The concentrations of TG were measured using a TG assay kit (Infinity Triglycerides kit) following the manufacturer’s protocol. The colormetric readings were collected with a microplate reader at OD 570 nm and the data were normalized to total proteins.

### RNA preparation and real-time PCR

Isolation of total RNA and reverse transcription into single-stranded cDNA was performed as previously described^70^. Gene expression was assessed in at least triplicate per sample by real-time quantitative PCR using the StepOnePlus Real-Time PCR System (Thermo Fisher Scientific) and SYBR Green chemistry with gene-specific primers at a final concentration of 200 nM. Primer sequences are provided in the Supplemental Materials (**Supplemental Table 1**). Relative expression levels were calculated using the 2-ΔΔCt method, with acidic ribosomal phosphoprotein P0 (RPLP0) for rat samples or hypoxanthine guanine phosphoribosyl transferase 1 (Hprt) for mouse samples as the housekeeping gene. Each experiment was independently repeated a minimum of three times. Gene expression values were normalized to the housekeeping genes included in the array and reported as fold change relative to control (CTL) mice or vehicle-treated cells.

### siRNA and plasmid transfection

siRNAs against rat *Cul1* (5’-AGCGCAGTTACCAACGGTTTA-3’), *Cul2* (5’TTCGAGCGACCAGTAACCTTA-3’), *Cul4b* (5’-ATGGGTTATTGGCCAACATAT-3’), *Cul9* (5’-CCGACGGACTTGTCTCTTCTA-3’) and luciferase (5’-AACGTACGCGGAATACTTCGA-3’) were used. Briefly, NRVCs were transfected with siRNAs (100 pmol per 2x10^6^ cells) using Lipofectamine RNAimax (Thermo Fisher Scientific) following the manufacturer’s protocol at 24∼48 hours after plating. Six hours after the transfection, the siRNA-containing medium was replaced with fresh medium containing 2% FBS. In some experiments, a second round of siRNA transfection may be performed 2 days after the first transfection to achieve sustained gene silencing.

## Statistical Analysis

All statistical analyses were performed using appropriate tests based on experimental design and data structure. Comparisons among more than two groups were analyzed using one-way ANOVA followed by the appropriate post hoc multiple-comparison tests. For experiments involving comparisons between two groups, an unpaired Student’s *t*test was used. When datasets included multiple technical or biological repeats nested within experimental conditions, a nested *t*-test was applied to account for intra-group variability. Survival curves were analyzed using standard survival analysis methods, including log-rank (Mantel-Cox) testing. Details of the specific statistical test applied to each dataset are indicated in the corresponding figure legends. Unless otherwise stated, data are presented as mean ± SEM, and statistical significance was defined as *, 0.05 > p > 0.01; **, 0.01 > p 0.001; ***, 0.001> p > 0.0001; ****, 0.0001 > p.

## ACKNOWLEDGEMENTS

This study was in part supported by the US National Institutes of Health grants (1R01HL173796-01, 1R01HL165205, R01HL124248-06A1, 1R01HL175569, and 1R01HL175569 to H.S. and R01HL146807A1 to J.L.) and the American Heart Association grant (959479 to H.S. and 19CDA34760311 and 17POST33661272 to J.Z.).

## Notes

### Competing Interest Statement

The authors have declared no competing interest.

