## Supplemental Figures for "Neddylation Regulates Mitochondrial Dynamics and Turnover in the Adult Heart"

### Supplimental Figure 1

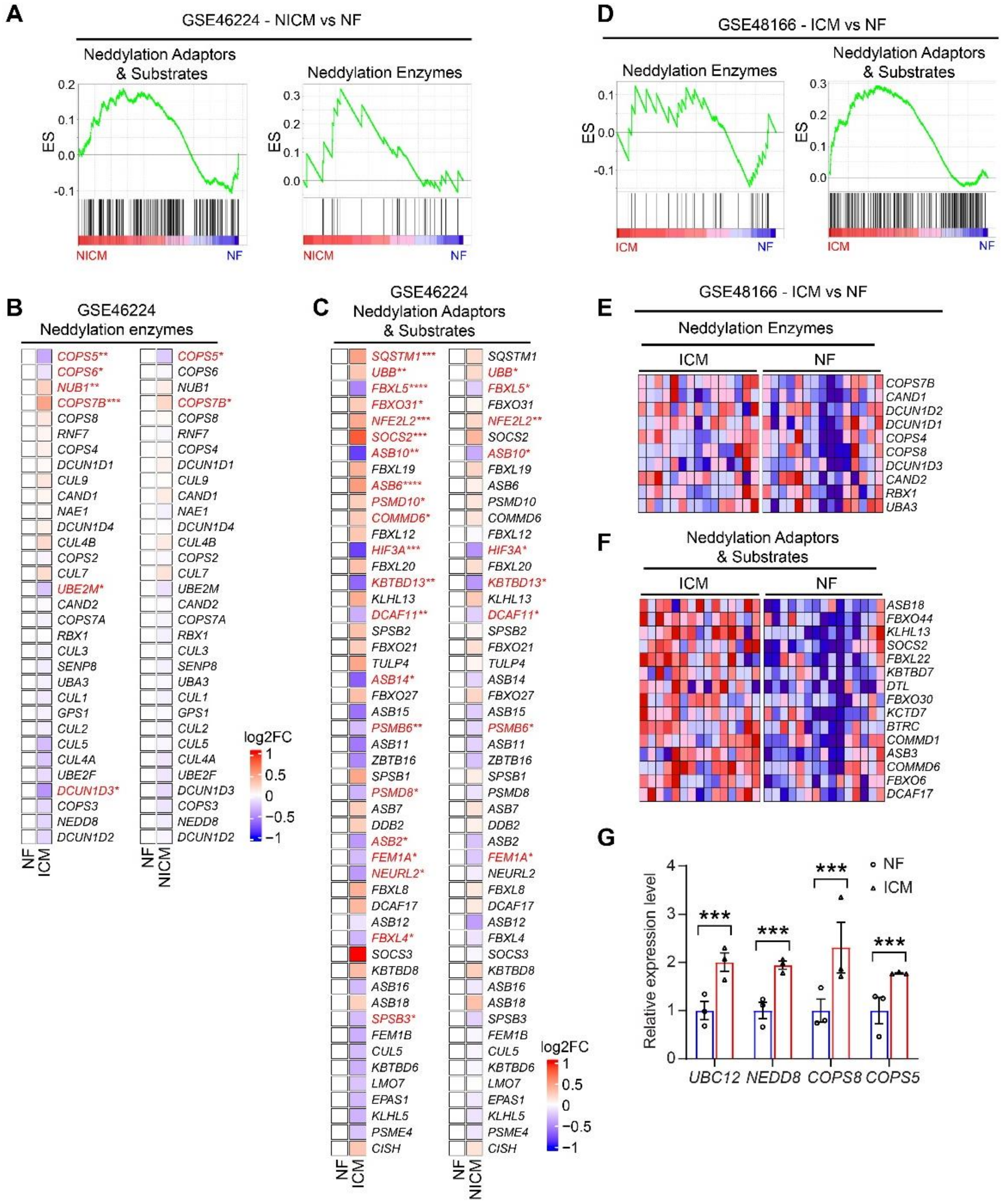


**Supplemental Figure 1.** Enrichment analyses of neddylation pathways for indicated sample sets. ICM, ischemic cardiomyopathy; NICM, non-ischemic cardiomyopathy; NF, non-failing heart. **A,** GSEA of patient samples in GSE46224 on indicated gene sets. **B-C,** Heatmap of expression level of indicated genes within the sets in panel **A** and **Fig. 1B**. **D,** GSEA of patient samples in GSE48166 on indicated gene sets. **E-F,** heatmap of expression level of indicated genes within the sets in panel **D**. **G,** qPCR analysis revealing expression level of indicated genes for human patient heart samples in **Fig. 1F&G**. ***, P<0.001.

### Supplimental Figure 2

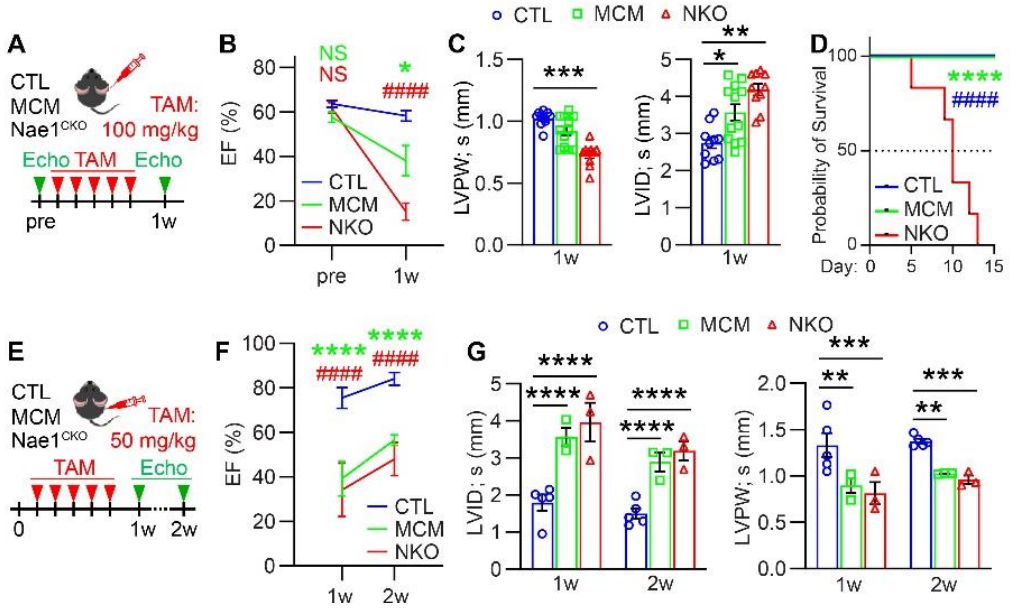


**Supplemental Figure 2. A,** schema of Nae1$^{CKO}$ mice injecting tamoxifen (TAM) at indicated dose (100 mg/kg) and time. **B**, echocardiography showing the ejection fraction (EF) at indicated time point in **A.** Red fonts indicate comparison of Nae1$^{CKO}$ (NKO) vs control (CTL) mice, green fonts indicate comparison of αMHCMer-Cre-Mer (MCM) vs CTL mice. **C**, echocardiography measurements of animals in **A** at 1 week post TAM injection. **D**, survival curve of animals shown in **A**. **E**, schema of Nae1$^{CKO}$ mice injecting tamoxifen (TAM) at indicated dose (50 mg/kg) and time. **F**, echocardiography showing the ejection fraction (EF) at indicated time point in **E.** Red fonts indicate comparison of Nae1$^{CKO}$ (NKO) vs control (CTL) mice, green fonts indicate comparison of αMHCMer-Cre-Mer (MCM) vs CTL mice. **G**, echocardiography measurements of animals in **E** at 1 and 2 weeks post TAM injection. LV, left ventricle; AW, anterior wall thickness; PW, posterior wall thickness; ID, internal diameter; s, systolic; d, diastolic. NS, not significant; *, 0.01<P< 0.05; **, 0.001<P< 0.01; ***, 0.0001<P< 0.001; ****, P< 0.0001; ####, P<0.0001.

**Supplimental Figure 3**

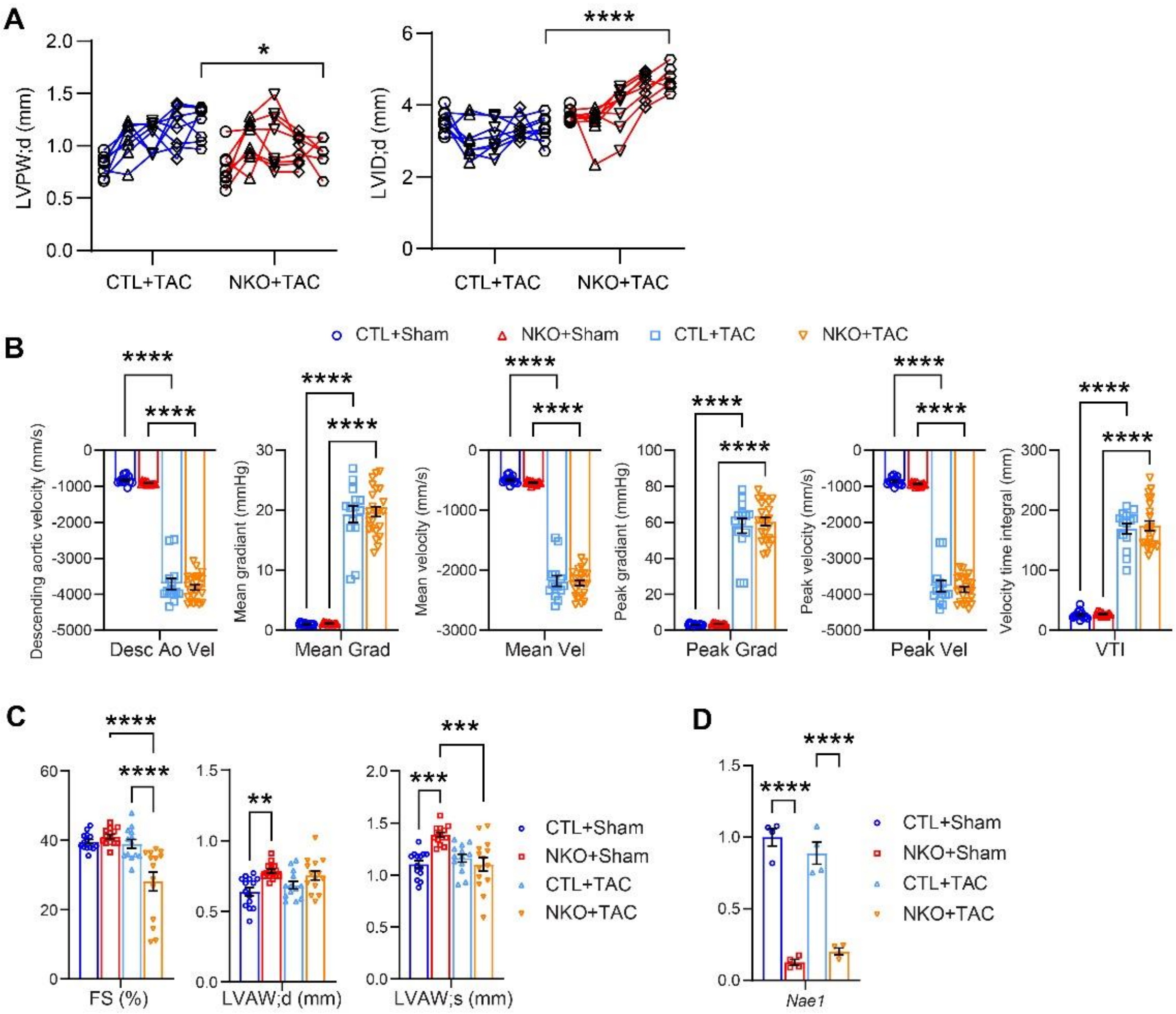


**Supplemental Figure 3. A,** time-trace echocardiography of Nae1$^{CKO}$ (KO) mice and control (CTL) mice after transverse aortic constriction (TAC) at pre-TAC, 1-, 2-, 3-, and 4-weeks post TAC. FS, fractional shortening; LVPW, left ventricle posterior wall thickness; LVAW, left ventricle anterior wall thickness; s, systolic. **B**, heart weight (Hw) body weight (Bw) ratio and heart weight (Hw) tibia length (TL) ratio of indicated animal hearts. **C**, echocardiography of Nae1$^{CKO}$ (KO) mice and control (CTL) mice after TAC or sham at 2 weeks post TAC. **D**, qPCR analyses of indicated genes. Ns, not significant; *, 0.01<P<0.05; **, 0.001<P<0.01; ***, 0.0001<P<0.001; ****, P<0.0001.

**Supplimental Figure 4**

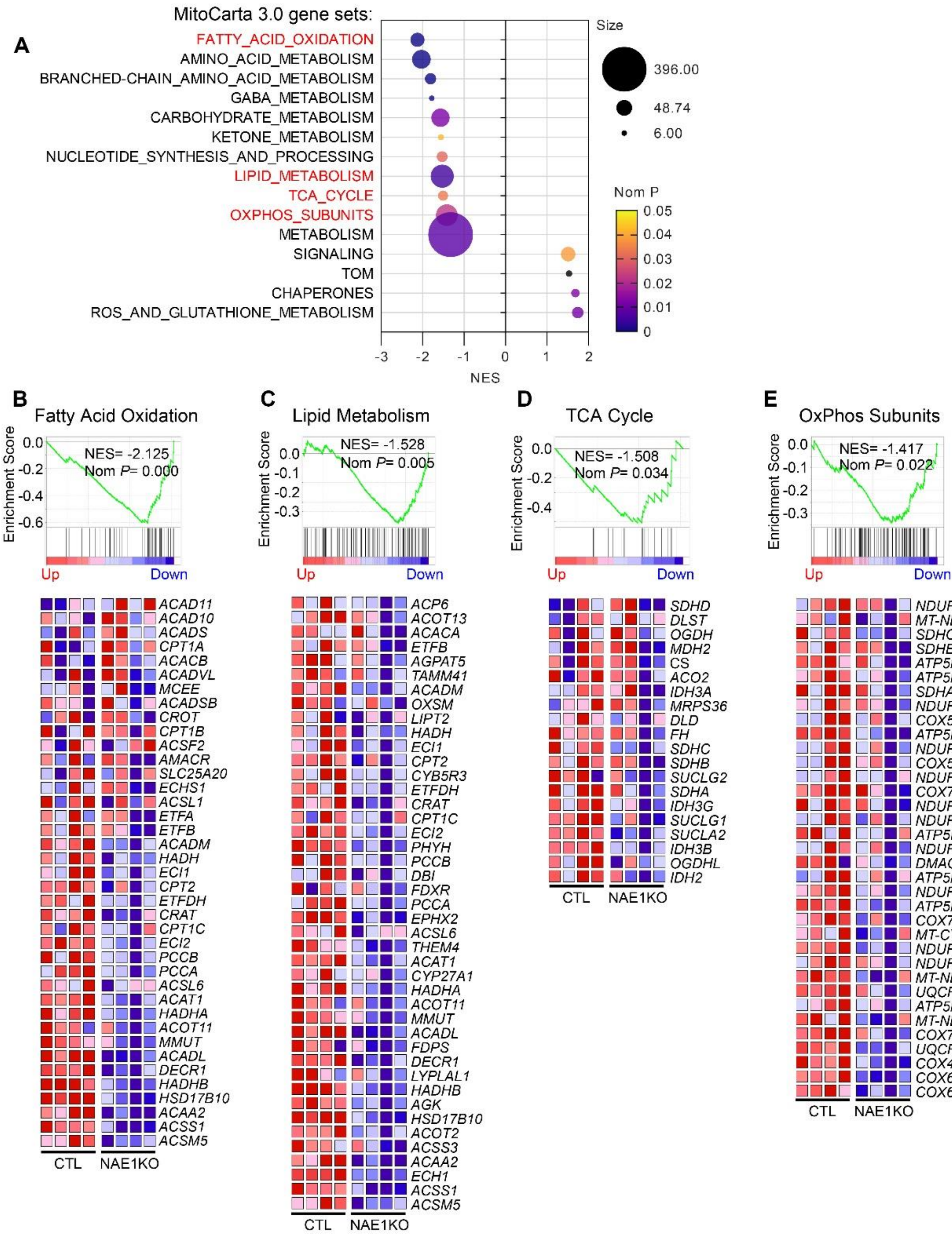


**Supplemental Figure 4. A,** enrichment analyses of MitoCarta gene sets for Nae1$^{CKO}$. **B-E,** GSEA of Nae1$^{CKO}$ (NAE1KO) mouse hearts vs control (CTL) (n= 4 vs 4) for fatty acid oxidation gene set (**B**), lipid metabolism gene set (**C**), TCA cycle gene set (**D**), and OxPhos subunit gene set (**E**), as well as the heatmap showing expression level of indicated genes within respective gene sets. NES, normalized enrichment score; Nom P, nominal P-value.

### Supplimental Figure 5

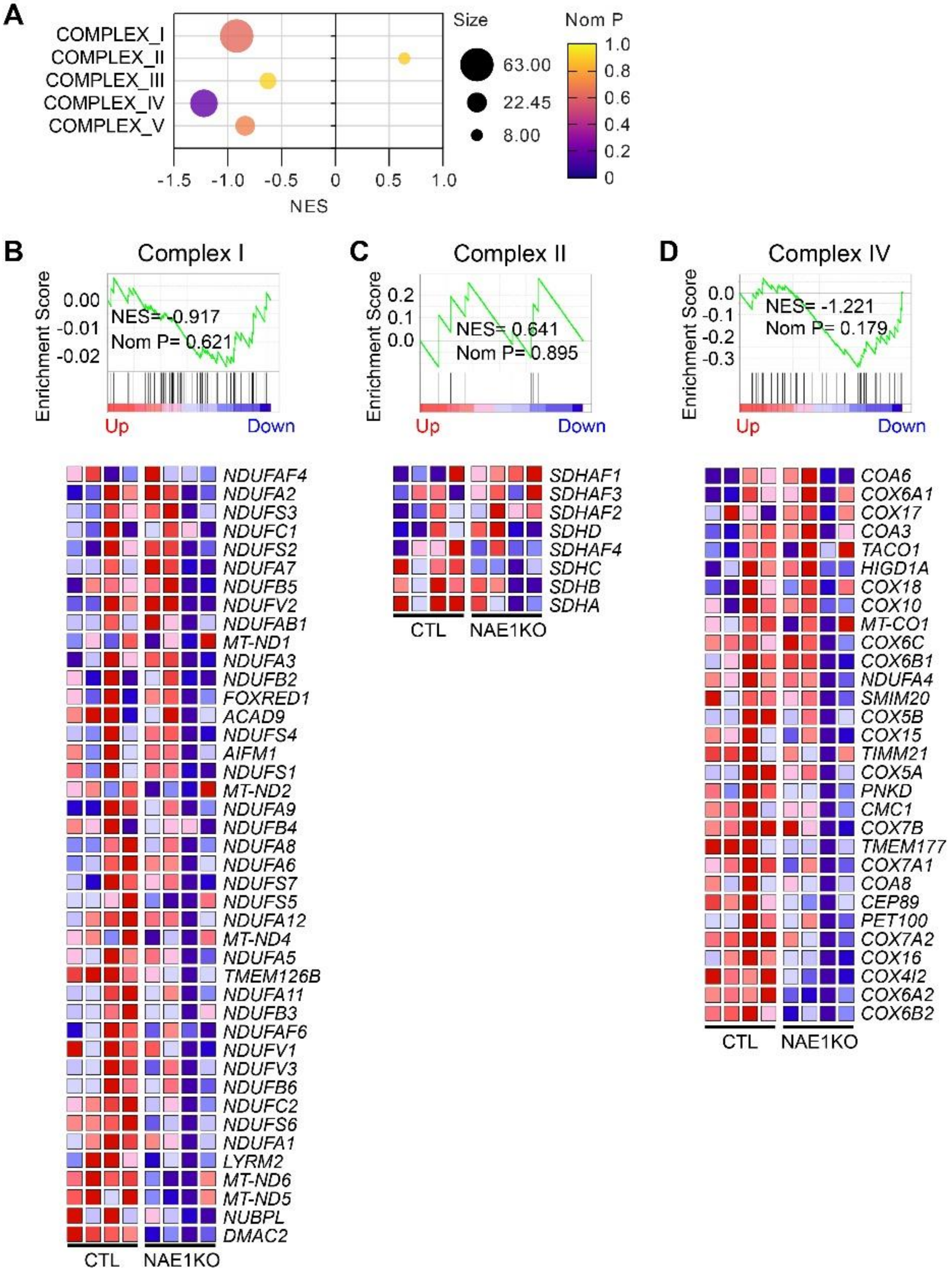


**Supplemental Figure 5. A,** dot plot indicating the NES, gene set size, and Nom P values of indicated gene sets. **B-D,** GSEA of Nae1$^{CKO}$ (NAE1KO) mouse hearts vs control (CTL) (n= 4 vs

4) for MitoCarta complex I (A), complex II (B), and complex V (C) gene sets, as well as the heatmap showing expression level of indicated genes within respective gene sets. NES, normalized enrichment score; Nom P, nominal P-value.

### Supplimental Figure 6

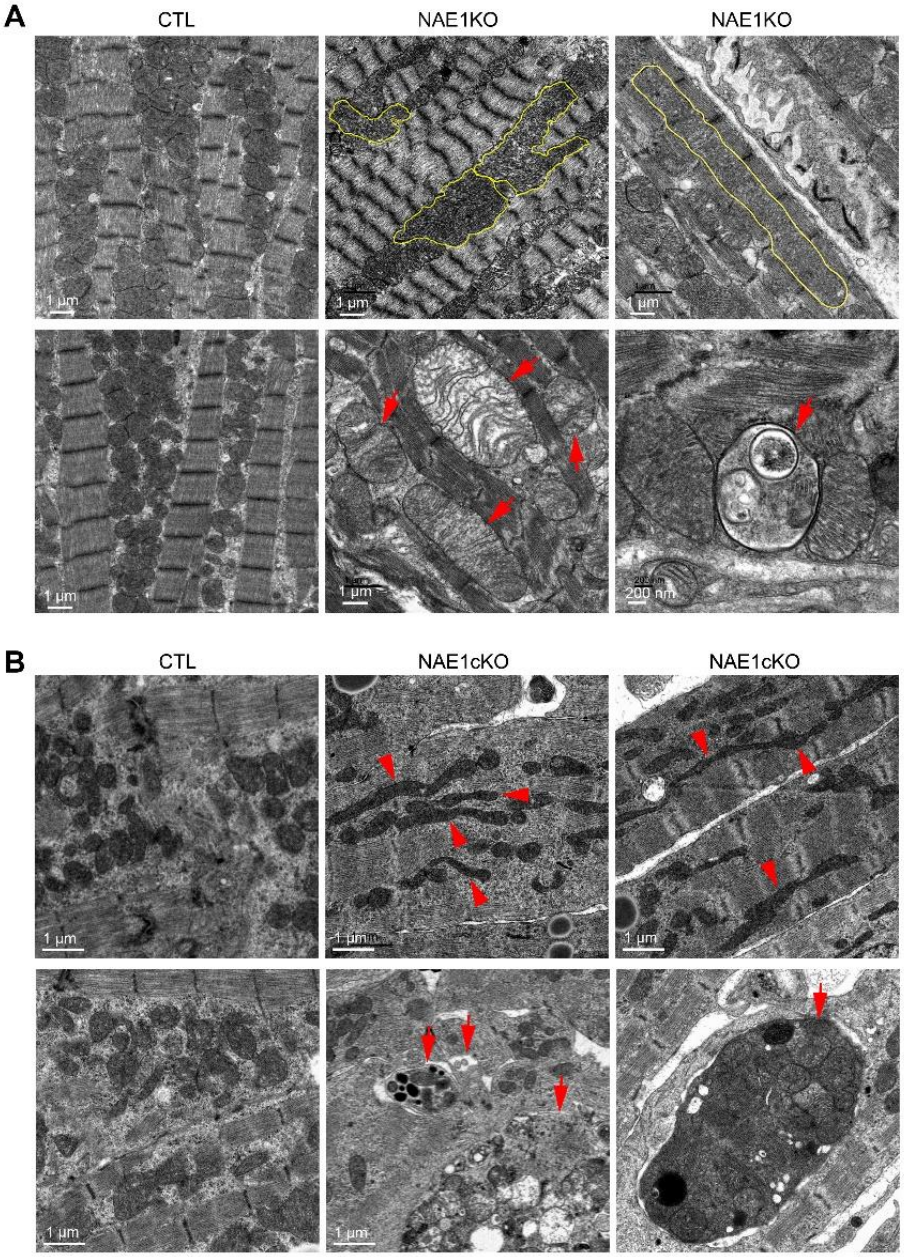


**Supplemental Figure 6. A,** TEM images of NAE1KO vs control hearts. **B**, TEM images of NAE1cKO vs control neonatal hearts. Yellow circle outlined elongated mitochondria in NAE1KO. Red arrowhead indicates elongated mitochondria in NAE1cKO hearts. Red arrow indicates mitochondrial remodeling or mitophagic vesicles.

**Supplimental Figure 7**

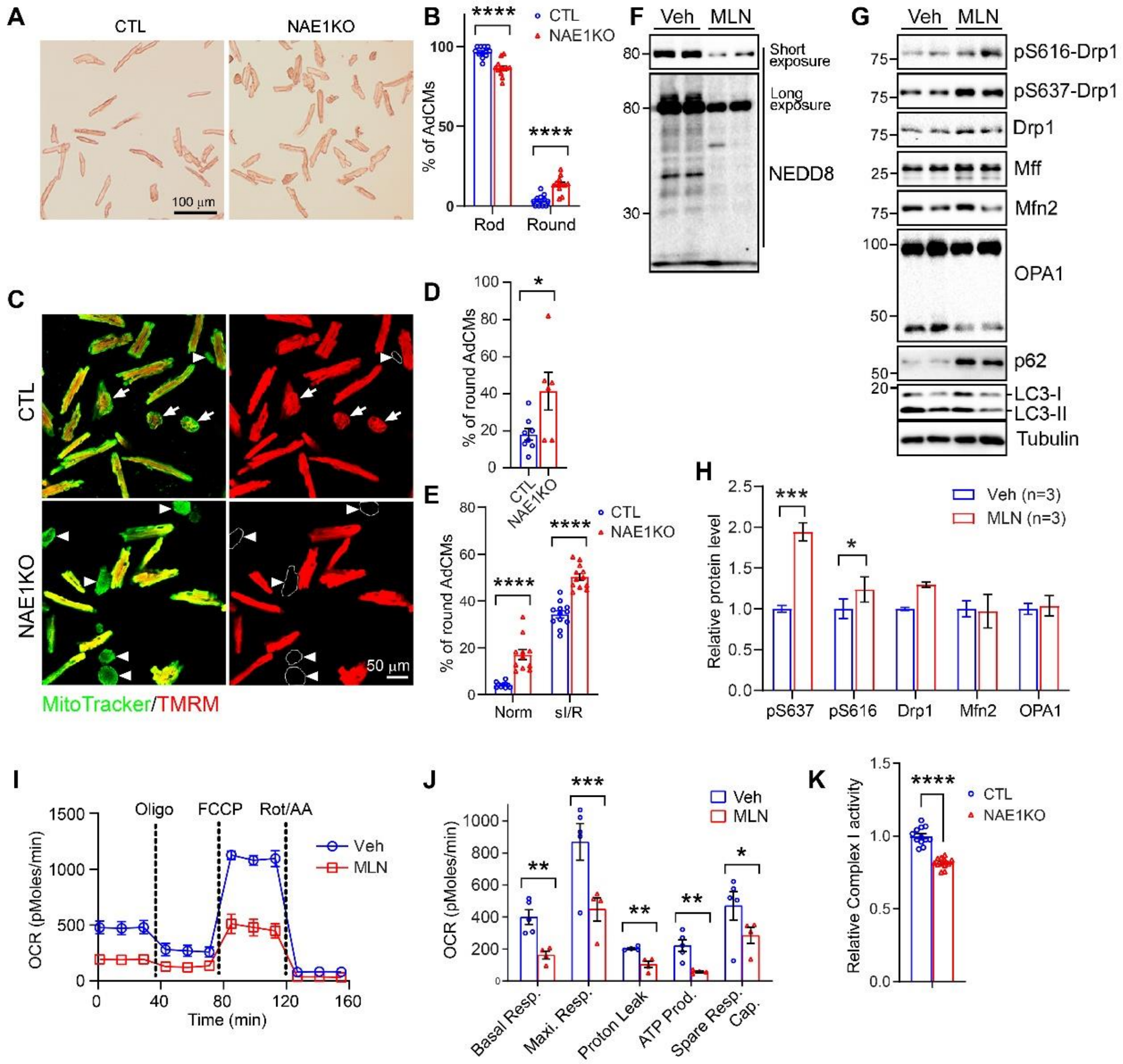


**Supplemental Figure 7. A**, bright field imaging of NAE1KO vs control (CTL) isolated adult cardiomyocytes (AdCMs). **B**, quantification of percentage of rod shaped AdCMs and round shaped AdCMs within indicated groups from panel **A**. **C**, live cell confocal fluorescent imaging of isolated AdCMs within indicated groups. Lyso, lysotracker; Mito, mitotracker. **D**, quantification of percentage of round AdCMs. **E**, quantification of percentage of round AdCMs treated under condition of normoxia or simulated ischemia-reperfusion. **F-G**, western blot showing neddylated proteins (**F**) or indicated protein levels (**G**) in MLN4924 (MLN) or vehicle (Veh) treated AdCMs isolated from wild type mice. **H,** quantification of **G**. **I-J,** seahorse Mito-stress assay and its quantifications within AdCMs treated with Vehicle or MLN for 72 hours. **K,** mitochondrial Complex I activity assay with freshly isolated mitochondria from mouse hearts as indicated. *, 0.01<P<0.05; **, 0.001<P<0.01; ***, 0.0001<P<0.001; ****, P<0.0001.

**Supplimental Figure 8**

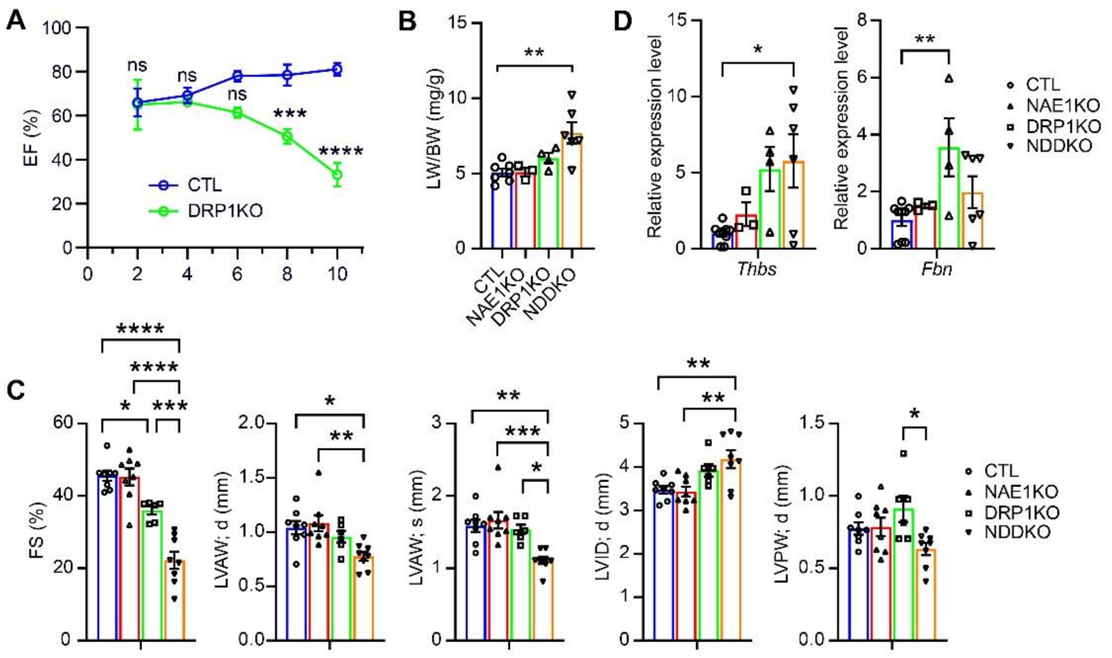


**Supplemental Figure 8. A**, time course echocardiography showing ejection fraction (EF) tracing tamoxifen-inducible αMHC$^{MCM}$-driven Drp1$^{CKO}$ (DRP1KO) vs control (CTL) mice after indicated weeks post tamoxifen injection. **B**, lung weight (LW) body weight (BW) ratio of Nae1$^{CKO}$ (NAE1KO), Drp1$^{CKO}$ (DRP1KO), and Nae1-Drp1 double knockout (NDDKO) mice upon collection at 6 weeks post TAM. **C**, echocardiography of indicated mice at 4 weeks post TAM. LV, left ventricle; AW, anterior wall thickness; PW, posterior wall thickness; ID, internal diameter; s, systolic; d, diastolic. **D**, qPCR analyses of indicated genes in heart tissue collected from indicated animals. Ns, not significant; *, 0.01<P<0.05; **, 0.001<P<0.01; ***, 0.0001<P<0.001; ****, P<0.0001.

**Supplimental Figure 9**

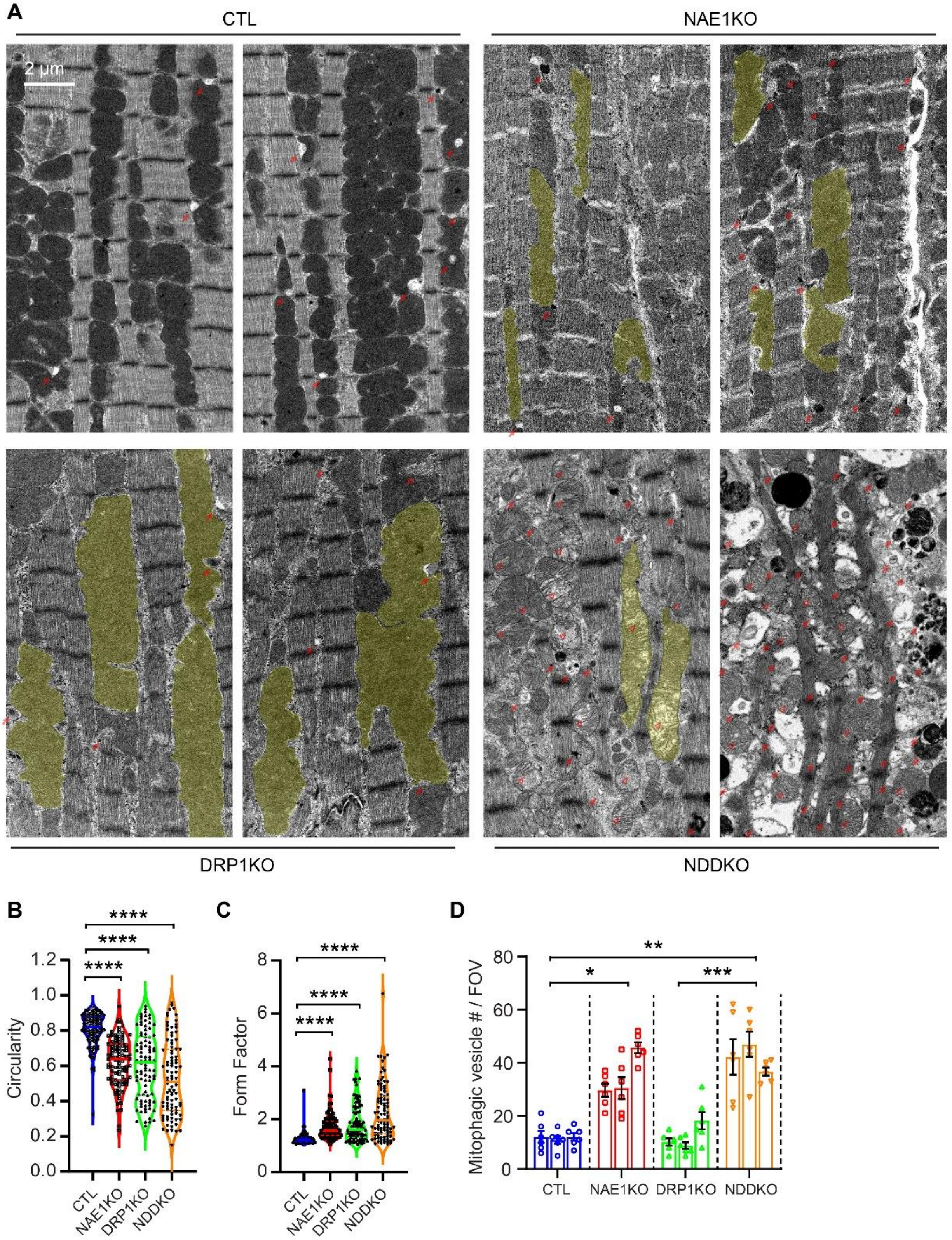


**Supplemental Figure 9. A**, transmission electron microscopy (TEM) of heart tissue from control (CTL), Nae1$^{CKO}$ (NAE1KO), Drp1$^{CKO}$ (DRP1KO), and Nae1-Drp1 double knockout (NDDKO)

mice. Solid red arrow indicates mitophagic vesicles; hollowed red arrowhead indicates mitochondrial remodeling; yellow shade indicates elongated (hyperfused) mitochondria. **B-D**, quantification of circularity of mitochondria (**B**), form factor (**C**), and mitophagic vesicle number per field of view (**D**) within indicated groups. Ns, not significant; *, 0.01<P<0.05; **, 0.001<P<0.01; ***, 0.0001<P<0.001; ****, P<0.0001.

**Supplimental Figure 10**

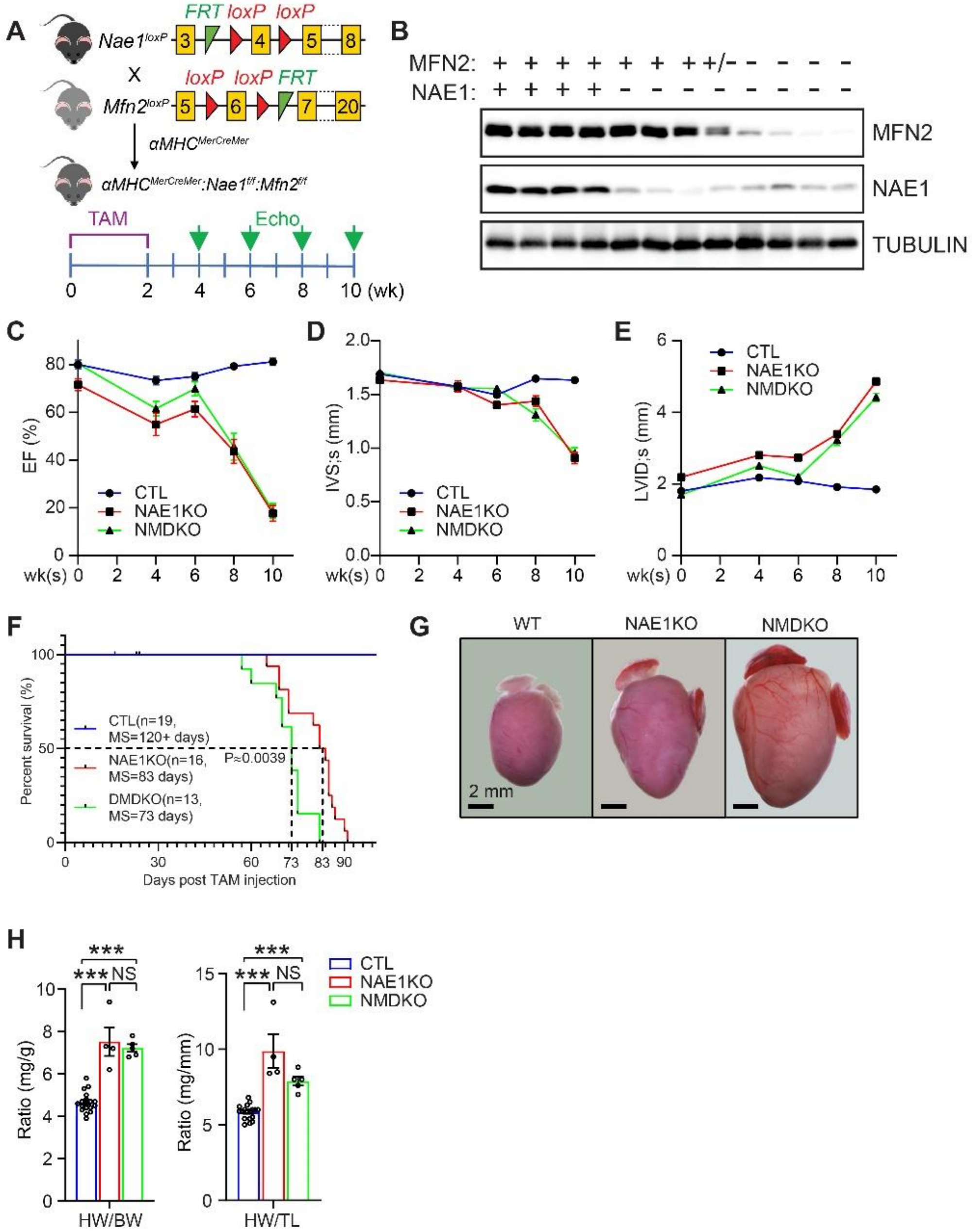


**Supplemental Figure 10.** **A**, scheme of the generation of *αMHC*$^{MerCreMer}$*:Nae1*$^{f/f}$*:Mfn2*$^{f/f}$ (*Nae1* and *Mfn2* double knockout, NMDKO). **B**, western blot (WB) of indicated proteins collected from mice of indicated genotypes 6 weeks after tamoxifen (TAM) injection. **C**, **D** & **E**, temporal echocardiography measurements of indicated parameters at 0, 4, 6, 8 and 10 weeks after TAM injection. EF, ejection fraction; LVID, left ventricle internal diameter; IVS, interventricular septum; s, systolic. **F**, survival curve of indicated mice after tamoxifen injection. **G**, gross morphology of indicated hearts collected at 10 weeks post TAM injection. **H**, heart weight/ body weight (HW/BW) and heart weight/ tibia length (HW/TL) ratio of hearts in indicated mice. Scale bar in panel **G**

represents 2mm as indicated. Statistical tests performed in panel **C**, **D** and **E**, were performed by two-way ANOVA followed by post hoc Tukey's multiple comparisons, and in panel **H** were performed by one-way ANOVA followed by Tukey's multiple comparisons, and in panel **F** was log rank test. *, 0.01<P< 0.05; **, 0.001<P< 0.01; ***, 0.0001<P< 0.001; ****, P< 0.0001; NS, not significant.

### Supplimental Figure 11

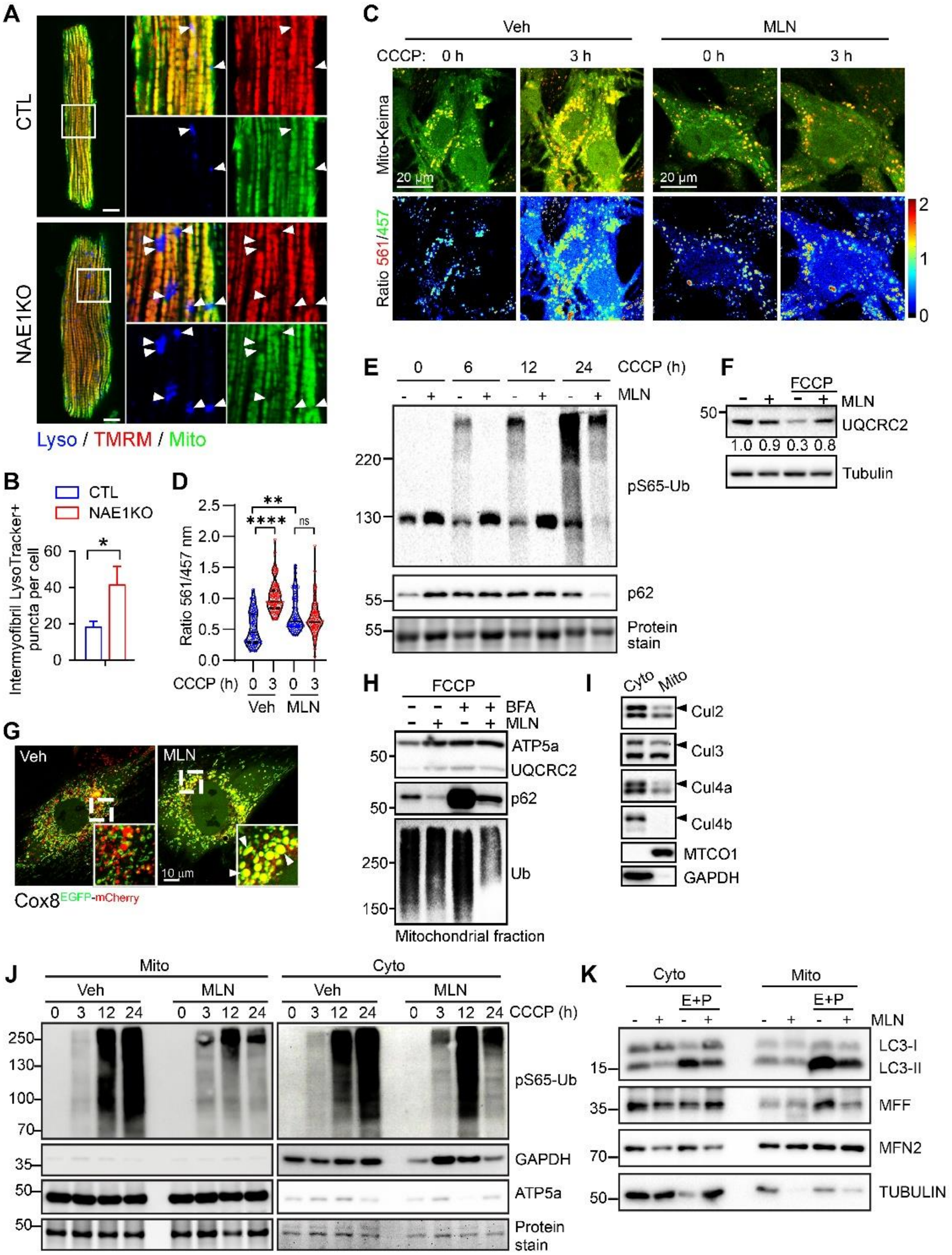

**Supplemental Figure 11. A,** live cell confocal imaging of isolated AdCMs within indicated groups. Lyso, lysotracker; Mito, mitotracker. **B**, quantification of number of lysotracker positive intermyofibril puncta staining per cell. **C**, NRVCs infected with Ad-Mito-Keima adenovirus and next subjected to MLN4924 (MLN, 1 μM, 48 hours) treatment, with or without CCCP (10 μM) treatment, and subjected to confocal microscopy for live cells. The ratio map was calculated by ratio of intensities within indicated channels and rendered in MATLAB as heatmap. **D**, quantification of ratios of indicated channels as represented in **C**. **E**, NRVCs treated with 1 μM MLN4924 (MLN) and with or without 10 μM CCCP for indicated time were blot with indicated antibodies. **F**, NRVCs treated with or without 1 μM MLN4924 (MLN) for 48 hours and with or without 10 μM CCCP for 24 hours (CCCP) were blot with indicated antibodies. **G**, NRVCs infected with Ad-COX8-EGFP-mCherry were subjected to 1 μM, 48 hours MLN or Vehicle and next observed under live cell confocal imaging. **H**, NRVCs treated or not with MLN4924 (1 μM, 48 hours), BFA (100 nM, 3 hours), CCCP (10 μM, 24 hours) were subjected to mitochondrial fractionation. The mitochondrial fractions were next analyzed by WB with indicated antibodies. **I**, 8-week-old adult mice hearts were subjected to mitochondrial fractionation and analyzed by WB with indicated antibodies. **J-K,** WB of NRVCs subjected to mitochondrial fractionation. Cyto, cytoplasmic fraction; Mito, mitochondrial fraction; E+P, E64d and Pepstatin A; Veh, vehicle control; MLN, 1 μM MLN4924 for 48 hours; CCCP, 10 μM for indicated time.

### Supplimental Figure 12

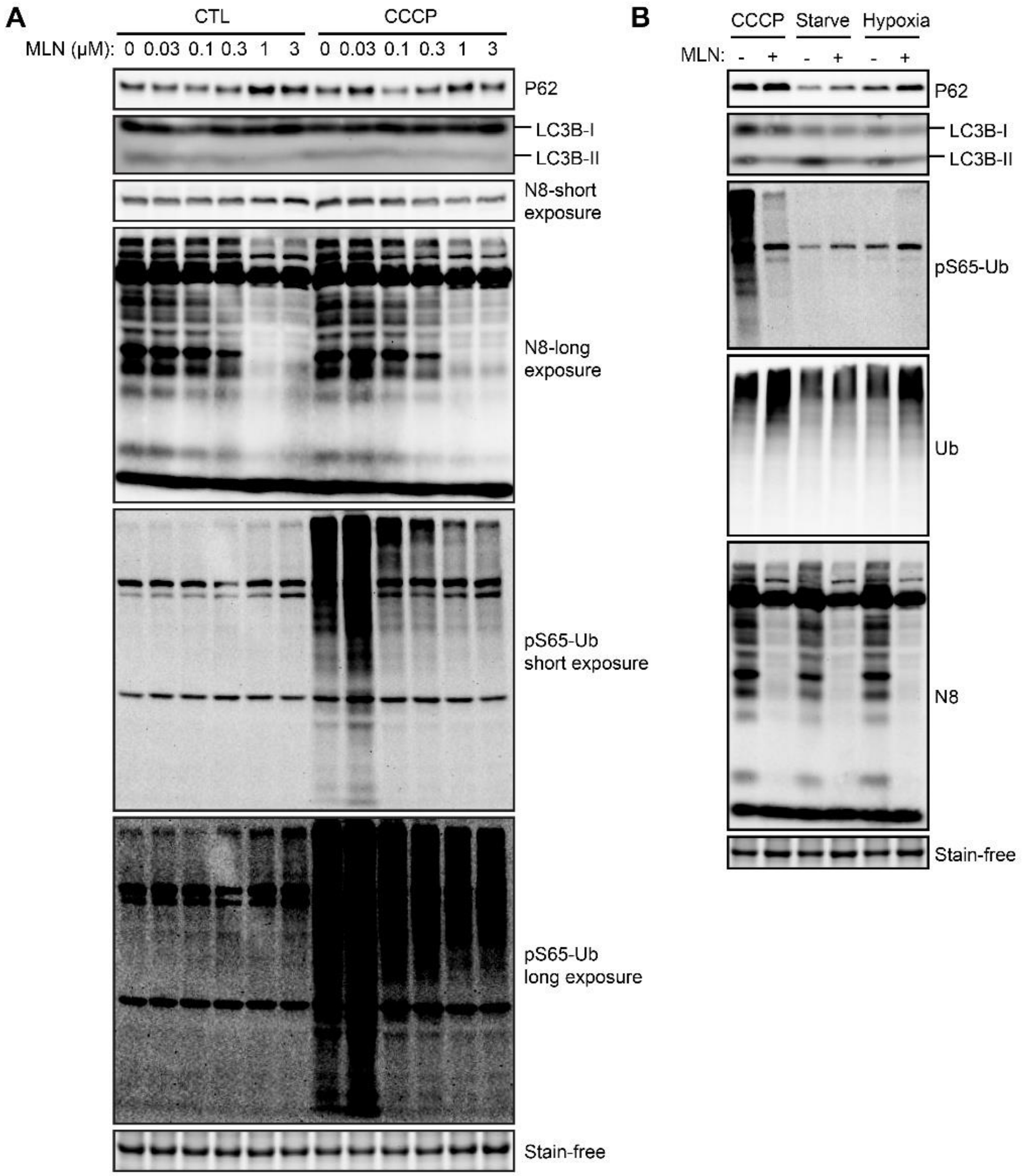


**Supplemental Figure 12.** *In vitro* western blot analyses within NRVCs with indicated antibodies. **A**, WB against antibodies as indicated of NRVCs treated with indicated concentration of MLN4924 for 48 hours, with or without 10 μM CCCP for 24 hours (CCCP). **B**, WB of NRVCs treated or not with 1 μM MLN for 48 hours, next subjected to 10 μM CCCP for 24 hours (CCCP), starved with 0% FBS media (Starve), or cultured in 1% $O_2$ hypoxia condition for 16 hours (Hypoxia).

### Supplimental Figure 13

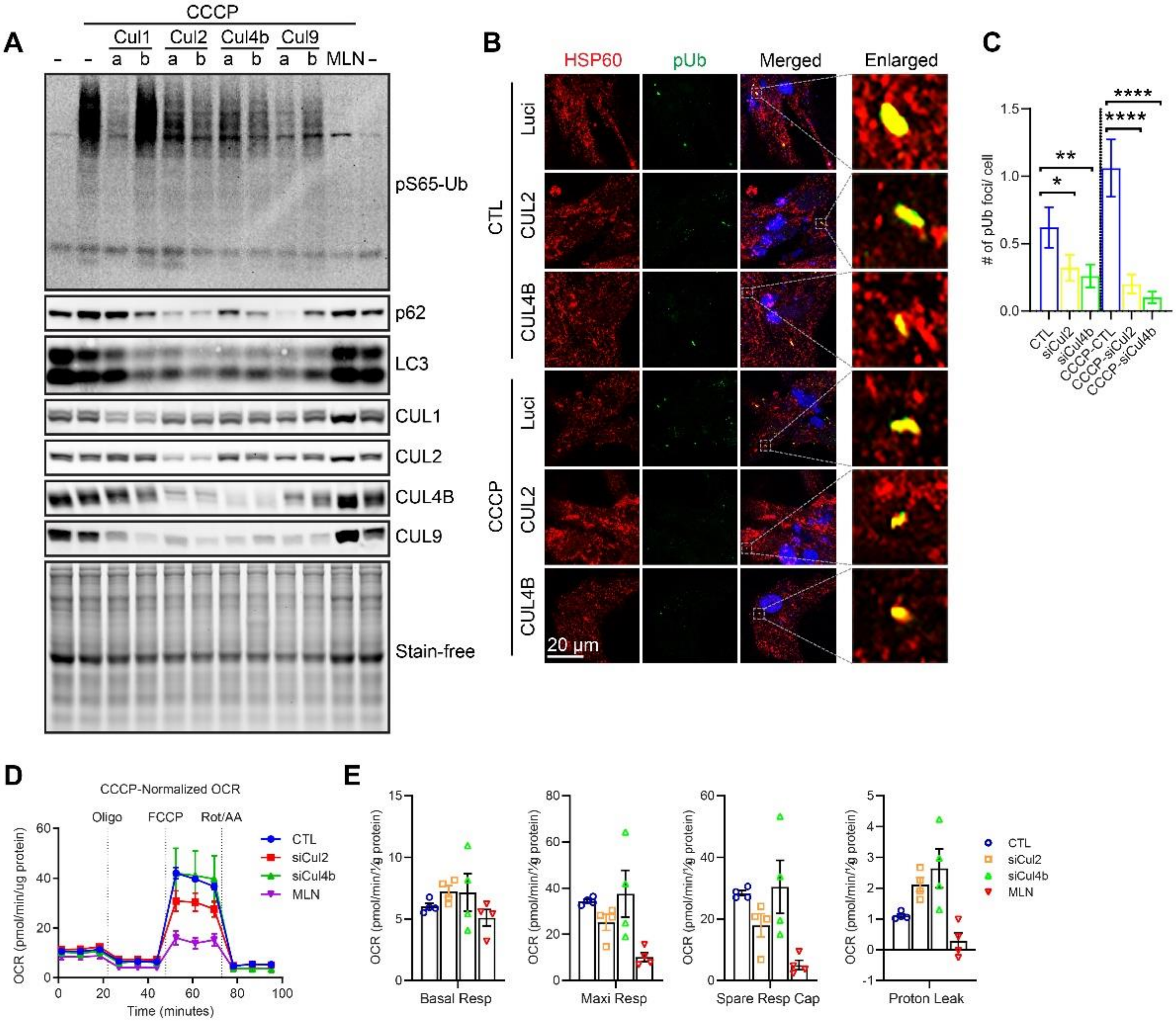


**Supplemental Figure 13. A,** NRVCs treated with siRNA against *Cul1*, *Cul2*, *Cul4b* and *Cul9* (a or b indicates siRNAs targeting different regions of the same gene) or Luciferase control (-), or treated with 1 μM MLN4924 for 48 hours, then treated with 10 μM CCCP for 24 hours or not, and subjected to western blot analyzing proteins as indicated. **B-C,** immunostaining of NRVCs treated with siRNAs against Luciferase control, Cul2, or Cul4b, followed by control or CCCP treatment (**B**) and the quantification of number of phospho-S65-Ub (pUb) foci per cell (**C**). **D-E**, seahorse analysis of Mito Stress assay on NRVCs with indicated treatments. *, 0.01<P< 0.05; **, 0.001<P< 0.01; ***, 0.0001<P< 0.001; ****, P< 0.0001.
