## Supplemental Tables for "Neddylation Regulates Mitochondrial Dynamics and Turnover in the Adult Heart"

### Supplemental Table 1

Antibodies:

| Proteins | Species | Vendor | Cat # | Dilution |
|---|---|---|---|---|
| CD45 | mouse | R&D Systems | AF114 | 1/50 (IF) |
| CUL1 | rabbit | Santa Cruz Biotechnology | sc-11384 | 1/1000 |
| CUL2 | rabbit | Thermo Fisher Scientific | 51-1800 | 1/1000 |
| CUL9 | rabbit | Invitrogen | PA5-20277 | 1/1000 |
| Cullin 4a | rabbit | Novus Biologicals | NB100-2267 | 1/1000 |
| DRP1 | rabbit | Cell signal | 5391 | 1/1000 |
| GAPDH | mouse | Santa Cruz Biotechnology | sc-32233 | 1/3000 |
| HSP60 | mouse | Thermofisher Scientific | MA3-012 | 1/50 (IF) |
| LAMP1 | rat | DSHB | 1D4B-c | 1/50 (IF) |
| LC3 | mouse | MBL Int. Corp. | M186-3 | 1/1000 |
| MFF | rabbit | Cell signal | 84580S | 1/1000 |
| MFN2 | rabbit | Cell signal | 9482S | 1/1000 |
| NAE1 | rabbit | Cell Signaling Technology | 14321 | 1/1000 |
| NEDD8 | rabbit | Cell Signaling Technology | 2754 | 1/250 |
| OPA1 | rabbit | Cell signal | 80471S | 1/1000 |
| P62 | guinea pig | American Res. Products | GP62-C | 1/2000 |
| Phalloidin | 568-conj | Invitrogen | A12380 | 1/200 (IF) |
| Phospho-Ubiquitin (Ser65) | rabbit | Cell Signaling Technology | 62802S | 1/500 (WB); 1/50 (IF) |
| Phospho-DRP1 (Ser616) | rabbit | Cell signal | 3455S | 1/1000 |
| Phospho-DRP1 (Ser637) | rabbit | Cell signal | 4867S | 1/1000 |
| TOM20 | rabbit | Cell Signaling Technology | 42406 | 1/25 (IF) |
| Total OXPHOS Cocktail | mouse | Abcam | ab110413 | 1/2000 |
| UBC12 | rabbit | Epitomics | 3690-1 | 1/500 |
| WGA | 488-conj | Invitrogen | W11261 | 1/200 (IF) |
| β-Tubulin | mouse | DSHB | E7 | 1/2000 |

Primers for qPCR:

| Gene | Species | Sequence (sense) | Sequence (antisense) |
|---|---|---|---|
| *UBC12* | human | GCGGATCCAGAAGGACATAAA | CACTCTTGTAGAAGCCCTCATC |
| *NEDD8* | human | GAGGATCCCAGGATTCAGTATTC | CAACCAGGGACACAGTCATAA |
| *CSN8* | human | AGTACCCAGATGGTGTTCTTTC | CACTTACTGCCCTGGAACTATT |
| *CSN5* | human | CCATTTGTAGCAGTGGTGATTG | AGGAGGTTTGTAGCCCTTTG |
| *Nae1* | mouse | CAACTCAGATCCCAAGCAGTAT | CCTTTAAGGCACGAGCTAGAA |
| *Myh6* | mouse | CGCATCAAGGAGCTCACC | CCTGCAGCCGCATTAAGT |
| *Myh7* | mouse | CGCATCAAGGAGCTCACC | CTGCAGCCGCAGTAGGTT |
| *Nppa* | mouse | CACAGATCTGATGGATTTCAAGA | CCTCATCTTCTACCGGCATC |
| *Nppb* | mouse | GTCAGTCGTTTGGGCTGTAAC | AGACCCAGGCAGAGTCAGAA |
| *Acta1* | mouse | AATGAGCGTTTCCGTTGC | ATCCCCGCAGACTCCATAC |
| *Serca2a* | mouse | TCGACCAGTCAATTCTTACAGG | CAGGGACAGGGTCAGTATGC |
| *Acaa2* | mouse | GACTTCTCTGCCACCGATTTA | TTGCCCACGATGACACTATC |
| *Acadvl* | mouse | CTTTGCAGGGACTCAAGGAA | CAAGCGAGCATACTGGGTATTA |
| *Dgat2* | mouse | GAAGGGCTTCTCTTCTCTTCAC | CTTTCTCCCAACGCCTCATAA |
| *Ech1* | mouse | CGCGATGACAGTTTCCAGTA | CAGAGATCGAAGGCTGATGTT |
| *Hadha* | mouse | AGACATCGGAGCTGTCTTTG | CACTACCTTCTGAGCACCATAC |
| *Mlycd* | mouse | CTGCCATCTTCTACTCCATCAG | GCTCCTTGACCACTCTCTTTAT |
| *Ppara* | mouse | AAGACTACCTGCTACCGAAATG | AACATTGGGCCGGTTAAGA |
| *Eci1* | mouse | CTGGACTTGCTGGAGATGTATG | AAGATCATGTTGGACGTGTAGAG |
| *Thbs* | mouse | CCTGGACTTGCTGTAGGTTATG | GTCATCATCTCTCTCGGTGTTG |
| *Fbn1* | mouse | CAGGCTCTTCTGTGTCGATATT | TGGCTGACAGCTACATTCATAG |
